# Distinct stimulatory mechanisms regulate the catalytic activity of Polycomb Repressive Complex 2 (PRC2)

**DOI:** 10.1101/210542

**Authors:** Chul-Hwan Lee, Marlene Holder, Daniel Grau, Ricardo Saldana-Meyer, Rais Ahmad Ganai, Jenny Zhang, Miao Wang, Marc-Werner Dobenecker, Danny Reinberg, Karim-Jean Armache

## Abstract

The maintenance of gene expression patterns during metazoan development is carried out, in part, by the actions of the Polycomb Repressive Complex 2 (PRC2). PRC2 catalyzes mono-, di-and trimethylation of histone H3 at lysine 27 (H3K27), with H3K27me2/3 being strongly associated with silenced genes. We demonstrate that EZH1 and EZH2, the two mutually exclusive catalytic subunits of PRC2, are differentially activated by various mechanisms. While both PRC2-EZH1 and PRC2-EZH2 are able to catalyze monomethylation, only PRC2-EZH2 is strongly activated by allosteric modulators and specific chromatin substrates to catalyze di-and trimethylation of H3K27. However, we also show that a PRC2 associated protein, AEBP2, can stimulate the activity of both complexes through a mechanism independent of and additive to allosteric activation. These results have strong implications regarding the cellular requirements for and accompanying adjustments in PRC2 activity, given the difference in the expression of EZH1 and EZH2 upon cellular differentiation.

## Introduction

Polycomb group (PcG) proteins are key epigenetic regulators that maintain transcriptional repression of lineage-specific genes throughout metazoan development, thereby contributing to the integrity of cell identity (Boyer et al., 2006; Buganim et al., 2013; Liang and Zhang, 2013; Margueron and Reinberg, 2011; Pasini et al., 2004; Pereira et al., 2010; Shen et al., 2008). In particular, PRC2 is responsible for the methylation of lysine 27 within histone H3 (H3K27me), with H3K27me3 being a hallmark of facultative heterochromatin (Margueron and Reinberg, 2011). PRC2 consists of three core subunits: one of two isoforms of Enhancer of zeste (EZH1 and-2), Embryonic ectoderm development (EED), and Supressor of zeste 12 (SUZ12) (Margueron et al., 2009; Murzina et al., 2008; Pasini et al., 2004; Tie et al., 2007). PRC2 core subunits are associated with a histone H4 binding protein: Retinoblastoma-associated proteins 46 or 48 (RbAp46/48) (Margueron and Reinberg, 2011; Murzina et al., 2008). The EZH1/2 subunit contains a SET domain that possesses histone methyltransferase (HMT) activity, and functions only when in complex with EED and SUZ12 (Nekrasov et al., 2005).

The catalytic activity of PRC2 is regulated by many factors including allosteric activators, incorporation of its different catalytic subunits (EZH1,-2), interactions with various histone modifications or chromatin structures, and PRC2 interacting partners including DNA and RNA (Beltran et al., 2016; Casanova et al., 2011; Davidovich et al., 2015; Holoch and Margueron, 2017; Jiao and Liu, 2015; Justin et al., 2016; Kaneko et al., 2014a, 2014b; Kim et al., 2009; Li et al., 2010, 2017; Liefke and Shi, 2015; Margueron et al., 2008; Mendenhall et al., 2010; da Rocha et al., 2014; Sanulli et al., 2015; Sarma et al., 2008a, 2014; Schmitges et al., 2011; Son et al., 2013; Yuan et al., 2012). The mechanism conveying allosteric activation of PRC2 was recently revealed by the crystal structures (Brooun et al., 2016; Jiao and Liu, 2015; Justin et al., 2016). The product of PRC2, H3K27me3, is recognized by an aromatic cage of its EED subunit, inducing a conformational change in PRC2 that specifically activates the EZH2 enzyme(Jiao and Liu, 2015; Justin et al., 2016). The hallmark of this mechanism entails the interaction between the Stimulatory Responsive Motif (SRM) of EZH2 and its SET-I domain (subdomain of SET) that stabilizes the SET domain (Jiao and Liu, 2015; Justin et al., 2016). The proposed model of this positive feedback loop involves: initial H3K27me3 deposition by PRC2, further PRC2 recruitment through binding of its EED subunit to H3K27me3 leading to allosteric activation of PRC2 and thus, additional H3K27me3 deposition giving rise to stable chromatin domains.

EZH1 and EZH2 are the PRC2 paralogs that contain the catalytic SET domain and are mutually exclusive when forming a complex with other PRC2 core subunits (Margueron et al., 2008; Shen et al., 2008). The catalytic activity of PRC2 containing EZH2 (PRC2-EZH2) is greater than that of PRC2 containing EZH1 (PRC2-EZH1)^11,12^. By contrast, PRC2-EZH1 possesses higher affinity to nucleosomes and can generate compacted chromatin structures independently from its catalytic function (Margueron et al., 2008; Son et al., 2013). Importantly, although PRC2-EZH2 can be allosterically activated by its own product, the reciprocal response in PRC2-EZH1 has not been well-demonstrated. Moreover, although the SET domain of EZH1 and EZH2 are 94% identical, only 65% identity is shared overall, suggesting that regions outside of the SET domain are responsible for the differences in their functional activity.

In addition to the canonical PRC2 core complexes, PRC2 forms additional complexes with various modulating cofactors including Jarid2, AEBP2, Polycomb-like proteins (PHF1, MTF2, and PHF19), EPOP, C10ORF12, and nucleic acids (DNA and RNA) (Alekseyenko et al., 2014; Beltran et al., 2016; Beringer et al., 2016; Grijzenhout et al., 2016; Kaneko et al., 2014b; Kim et al., 2009, 2011; Liefke and Shi, 2015; Liefke et al., 2016; Pasini et al., 2010; da Rocha et al., 2014; Sarma et al., 2014). Jarid2 is a well-known PRC2 interacting partner that enhances PRC2 activity by two independent manners. First, PRC2 trimethylates lysine 116 of Jarid2 (Jarid2-K116me3) and similarly to H3K27me3, allosterically activates PRC2 by binding to the aromatic cage of EED (Justin et al., 2016; Sanulli et al., 2015). Secondly, Jarid2 was shown to stimulate PRC2 activity by facilitating nucleosome binding (Son et al., 2013). Another PRC2 partner, AEBP2, is an evolutionarily conserved Gli-Krüppel (Cys2-His2)-type Zn finger protein that was shown to also stimulate the catalytic activity of PRC2 (Grijzenhout et al., 2016; Kaneko et al., 2014a). Currently, the underlying molecular mechanism by which AEBP2 stimulates PRC2 activity is not clear.

PRC2 is required to repress key differentiation genes in ESCs, and its components are highly regulated during differentiation. Once cells are terminally differentiated, the expression of EZH2 is significantly reduced in most cells and— dependent on the lineage commitment—specific PRC2 target genes are de-repressed (Boyer et al., 2006). In contrast to EZH2, EZH1 expression remains constant or is upregulated during differentiation (von Schimmelmann et al., 2016; Stojic et al., 2011). Knockout of EZH2 impairs cell differentiation due to unscheduled gene expression and deletion of both EZH1 and EZH2 results in more significant differentiation defects (Chou et al., 2011; Shen et al., 2008; Stojic et al., 2011). Moreover, an EZH2 null mutation results in lethality at early stages of mouse development, while EZH1 null mice are viable (Ezhkova et al., 2011; O’Carroll et al., 2001). Nonetheless, EZH1 has an important function during differentiation as well as in fully differentiated cells. For instance, EZH1 is required for hematopoietic stem cell maintenance, hair follicle homeostasis, and protection from neurodegeneration (Ezhkova et al., 2009, 2011; Hidalgo et al., 2012; von Schimmelmann et al., 2016). These results strongly suggest that the balance of PRC2-EZH1 and PRC2-EZH2 is critical for fine-tuning differentiation genes during development. In addition, expression of Jarid2 is significantly reduced during differentiation while AEBP2 remains generally constant (Hubbard et al., 2013; van de Leemput et al., 2014; Son et al., 2013), suggesting distinct roles for these cofactors in differentiated versus undifferentiated cells. To better understand the roles of PcG proteins during development, it will be necessary to determine the distinct functional roles of EZH1-and EZH2-containing PRC2 as well as other cofactors in the context of development.

PRC2 is also highly regulated by the context of chromatin (Martin et al., 2006; Schmitges et al., 2011; Yuan et al., 2012). The catalytic activity of PRC2-EZH2 is robustly increased on di/oligonucleosomes compared to mononucleosomes, suggesting that the simultaneous and multivalent interactions of PRC2 to two adjacent nucleosomes are required for optimal catalytic activity (Ciferri et al., 2012; Martin et al., 2006; Yuan et al., 2012). Moreover, chromatin that comprises shorter linker DNA, was reported to increase the HMT activity of PRC2-EZH2 (Yuan et al., 2012), however little is known regarding the impact of chromatin structure on the catalytic activity of PRC2-EZH1.

Here, we investigated the functional differences inherent to PRC2-EZH1 and PRC2-EZH2. Biochemical and genetic studies revealed marked differences in the catalytic activity of PRC2-containing these paralogues as a function of nucleosome repeat length and nucleosome-free DNA. We also provide new insights into the differential allosteric activation of PRC2-EZH1 and PRC2-EZH2. Furthermore, we demonstrate that these inherent differences result in differential sensitivity to both EED and SAM-based inhibitors of PRC2 enzymatic activity. Finally, we show that a basic region of AEBP2 is required for enhanced nucleosome binding and enhanced stimulation of both PRC2-EZH1 and-EZH2. Such AEBP2-mediated stimulation occurs through a mechanism distinct from that of allosteric activation by the methylated substrate.

## Results

### PRC2-EZH2 is the dominant H3K27 methyltransferase *in vitro and in vivo.*

To understand the distinct functions of EZH1 and EZH2, and their contributions to H3K27 methylation *in vivo*, we generated EZH1^−/−^(EZH1-KO), EZH2^−/−^(EZH2-KO), and EZH1^−/−^/EZH2-ΔSET (EZH1-KO/EZH2-ΔSET) in mESCs using the CRISPR-Cas9 approach (see Materials and Methods for details) and monitored H3K27 methylation via immunoblotting. As expected, the levels of H3K27me2/3 were significantly reduced in the EZH2-KO, while H3K27me1 was slightly increased, indicating that EZH2 contributes to H3K27me2 and-me3 while EZH1 was dispensable for higher order methylation states in mESCs (Figure 1A, lanes 1 and 2). Likewise, there was no effect on the levels of H3K27me1/2/3 in EZH1-KO lines when compared to WT (Figure 1A, lanes 3 and 4). These data suggested that EZH2 is also able to catalyze monomethylation on H3K27. Interestingly, all levels of H3K27me were depleted in EZH1-KO/EZH2-ΔSET (Figure 1A, lane 5) suggesting that EZH1 is an H3K27 monomethyltransferase. Together, these data indicate that the dominant H3K27 methyltransferase activity is derived from PRC2-EZH2 in mESCs. Moreover, EZH1 contributes to H3K27 monomethylation in mESCs, while EZH2 contributes to all states of H3K27 methylation. Finally, the data conclusively show that EZH2 and EZH1 are the only enzymes able to catalyze H3K27 monomethylation in mESCs.

**Figure 1.**
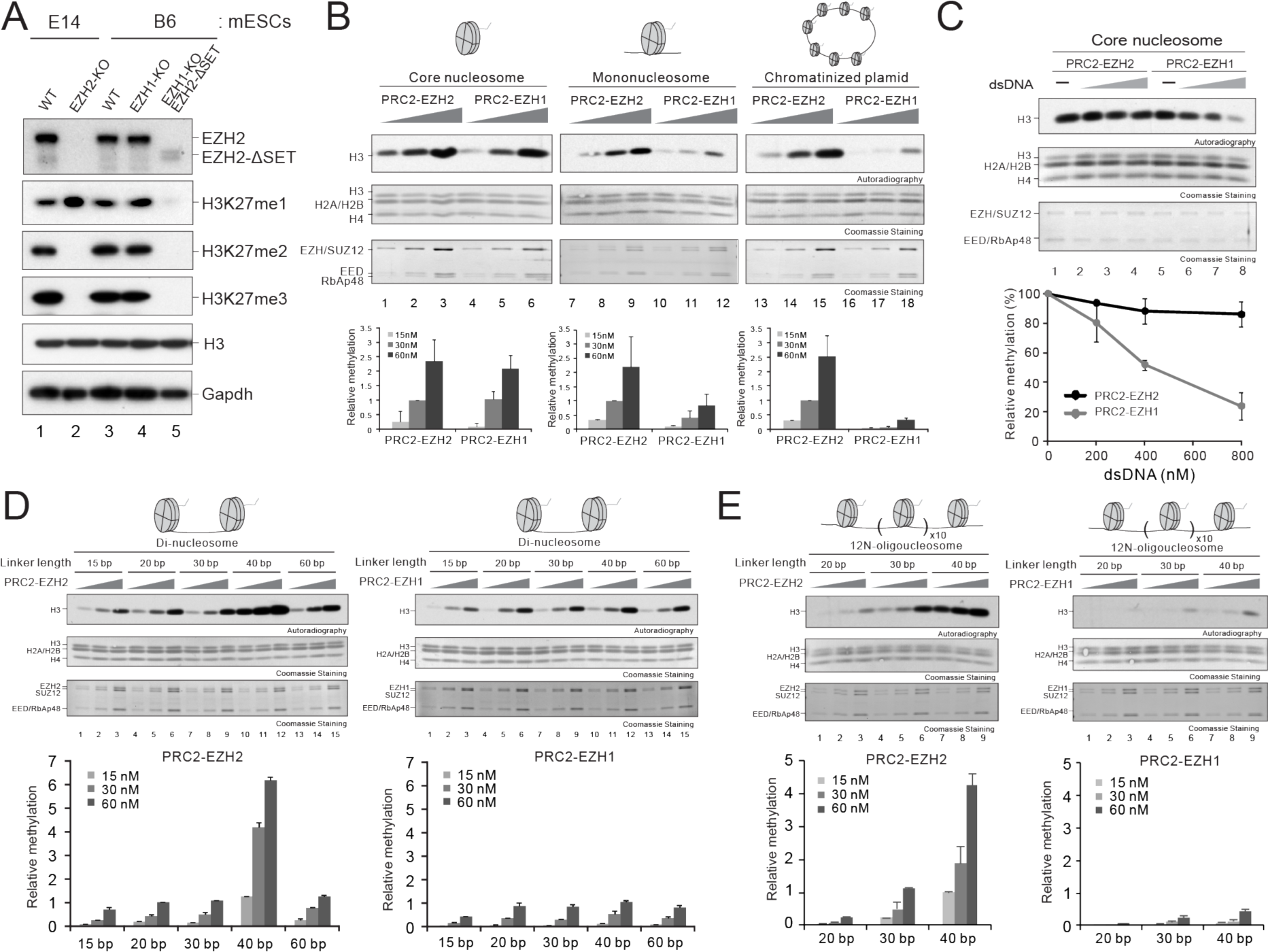
Substrate preference of PRC2-EZH1 and PRC2-EZH2 differentiate their intrinsic catalytic activity. (A) Western blot analysis for EZH2, H3K27me1, H3K27me2, H3K27me3, Gapdh and total histone H3 levels from E14 mESCs WT cells or cells carrying EZH2-KO, B6 mESCs WT cells or cells carrying EZH1-KO or EZH1-KO/EZH2ASET, as indicated. (B-E) Histone methyltransferase assays. The details of HMT assay conditions as well as the generation of the nucleosome substrates are described in the Materials & Methods. (B) *Top*, HMT assays containing PRC2-EZH1 or PRC2-EZH2 (15, 30, or 60 nM) using core nucleosomes, mononucleosomes, or chromatinized plasmids as substrates (300 nM) as illustrated on the panel. The levels of methylation on histone H3 are shown by autoradiography (*Top* image). Coomassie blue staining of SDS-PAGE gels containing nucleosomes *(Middle* image) or PRC2 components *(Bottom* image) was used to visualize the relative concentration of each component present in each reaction. *Bottom*, quantification of the relative amount of ^3^H-SAM incorporated into histone H3 after 60 minutes of incubation (n=3 for each data point). (C) *Top*, HMT assays containing PRC2-EZH1 or PRC2-EZH2 (15 nM) using core nucleosomes (300 nM) as the substrate with increasing amounts of 75 bp double-stranded (ds) DNA (200, 400, or 800 nM). Other panels are as described in (B). *Bottom*, quantification of the relative amount of ^3^H-SAM incorporated into histone H3 after 60 minutes of incubation (n=3 for each data point). (D) *Top*, HMT assays containing PRC2-EZH1 or PRC2-EZH2 (15, 30, or 60 nM) using di-nucleosomes (300 nM) containing different lengths of linker DNA (15, 20, 30, 40, or 60 bp) as substrates. Note that the flanking DNA at both ends is not present. Other panels are as described in (B). (E) *Top*, HMT assays containing PRC2-EZH1 or PRC2-EZH2 (15, 30, or 60 nM) using 12N oligo-nucleosomes (300 nM) containing different lengths of linker DNA (20, 30, or 40 bp) as substrates. Other panels are as described in (B).

The expression level of EZH2 is approximately four times higher than that of EZH1 in ESCs (Hubbard et al., 2013; van de Leemput et al., 2014), suggesting that the higher EZH2 activity could simply be due to the abundance of the protein. However, in differentiated cells with extremely reduced levels of EZH2, the levels of H3K27 di-trimethylation are still mainly dependent on the remaining EZH2 (Gonzalez et al., 2009; Mu et al., 2013; Son et al., 2013). Therefore, not only their expression levels, but also their intrinsic enzymatic properties must contribute to the distinctive behaviors of EZH1 and EZH2. Thus, we revisited the previous *in vitro* findings to explore the difference between PRC2-EZH1 and PRC2-EZH2 activities (Margueron et al., 2008; Shen et al., 2008; Son et al., 2013). We previously reported that recombinant PRC2-EZH2 exhibited greater levels of catalytic activity than PRC2-EZH1 (Margueron et al., 2008; Son et al., 2013), while other reports found no difference (Shen et al., 2008). We reasoned that these inconsistent results were likely due to the HMT assay conditions, especially with respect to the different types of nucleosome substrates used. To address these discrepancies, we purified recombinant human PRC2 complex (EZH, EED, SUZ12, and RbAp48) containing either EZH1 or EZH2 (Figure S1) and monitored the activity of PRC2-EZH2 and PRC2-EZH1 using various chromatin substrates, such as core nucleosomes (no flanking DNA), mononucleosomes (flanking DNA on both sides), and chromatinized plasmids.

Surprisingly, the activity of PRC2-EZH2 and PRC2-EZH1 was relatively comparable when using core nucleosomes as the substrate (Figure 1B, *Left).* However, the activity of PRC2-EZH1 was lower on mononucleosomes and significantly lower on chromatinized plasmids when compared to PRC2-EZH2 (Figure 1B, *Middle* and *Right).* These data suggested that in contrast to PRC2-EZH2, PRC2-EZH1 was sensitive to free DNA. To test this possibility, we compared the activity of each complex on core nucleosomes in the absence or presence of double-stranded DNA (dsDNA) as competitor. As hypothesized, the addition of increasing amounts of free DNA reduced the activity of PRC2-EZH1, while PRC2-EZH2 was less affected (Figure 1C). Additionally, we found that PRC2-EZH1 exhibited higher affinity for DNA relative to PRC2-EZH2 (Figure S2). Together, these data suggested that the presence of free DNA significantly inhibited the HMT activity of PRC2-EZH1.

### EZH2 catalytic activity is highly stimulated by specific nucleosome repeat lengths

Many histone modifiers—including PRC2—bind nucleosomes in a multivalent manner, and their activity is sensitive to the length of linker DNA (Ciferri et al., 2012; Huh et al., 2012; Lee et al., 2013; Martin et al., 2006). To compare such sensitivity in the case of PRC2-EZH1 and PRC2-EZH2, we designed di-nucleosomes with variable linker lengths (Figure S3). Importantly, these nucleosome substrates were designed to allow for the specific measurement of enzymatic activity as the function of linker DNA length as they lack flanking, ‘free’ DNA, which impacts PRC2-EZH1 activity (Figure 1D and S3). PRC2-EZH1 was almost unaffected by the length of the linker DNA, however, PRC2-EZH2 sensed the linker difference in a non-linear fashion (Figure 1D). The activity of PRC2-EZH2 increased as the length of the linker DNA was increased up to 40 bp, and then declined as the length was further increased (Figure 1D and S4). Interestingly, the activity of PRC2-EZH2 on 40 bp linker dinucleosomes was 5-6 fold higher than that on 30 bp linker dinucleosomes (Figure 1D, *Left*, compare lanes 7-9 with lanes 10-12). The 40 bp preference was also corroborated using 12N oligonucleosomes as substrates (Figure 1E). In accordance, PRC2-EZH2 activity was significantly increased on 12N oligonucleosomes with 40 bp linker DNA, while PRC2-EZH1 activity was only marginally increased (Figure 1E). Based on these results, we speculated that PRC2-EZH2 localizes in between adjacent nucleosomes *in vivo.* We analyzed MNase-seq and SUZ12 ChIP-seq data, and observed that the occupancy of nucleosomes was lower at the center of SUZ12 binding peaks than the flanking regions (Figure S5), confirming our hypothesis. Together, these results suggested that PRC2-EZH2 does bind between nucleosomes and that its activity is regulated by the length of linker DNA. More importantly, the HMT activity of PRC2-EZH2 was significantly enhanced just by adjusting the length of linker DNA, suggestive of an alternative stimulatory mechanism specific for PRC2-EZH2 (see below).

### The Stimulatory Recognition Motif (SRM) underlies differential allosteric activation of PRC2-EZH2 and PRC2-EZH1

Allosteric activation of PRC2 is a critical mechanism that enhances its HMT activity and is key to ensuring the optimal propagation of H3K27me3 (Justin et al., 2016; Margueron et al., 2009). Structural studies of PRC2 containing EZH2 revealed that its activity was significantly increased by H3K27me3 or Jarid2-K116me3 (Jiao and Liu, 2015; Justin et al., 2016). However, it is not known whether PRC2-EZH1 is also allosterically activated. Thus, we first monitored the activities of PRC2-EZH1 and PRC2-EZH2 in the absence or presence of an H3K27me3 peptide. Consistent with the results presented above (Figure 1B), the catalytic activities of PRC2-EZH1 and PRC2-EZH2 were similar on core nucleosomes in the absence of peptide (Figure 2A, lanes 1 and 5). Strikingly, the activity of PRC2-EZH2 was increased by approximately 12-fold in the presence of H3K27me3, while the activity of PRC2-EZH1 was only enhanced 2-fold (Figure 2A, *Left*, compare lanes 2-4 with lanes 6-8). The differential response was also observed when using chromatinized plasmids or dinucleosomes comprising different lengths of linker DNA as substrates (Figure 2A, *Right*, Figure S6, respectively).

**Figure 2.**
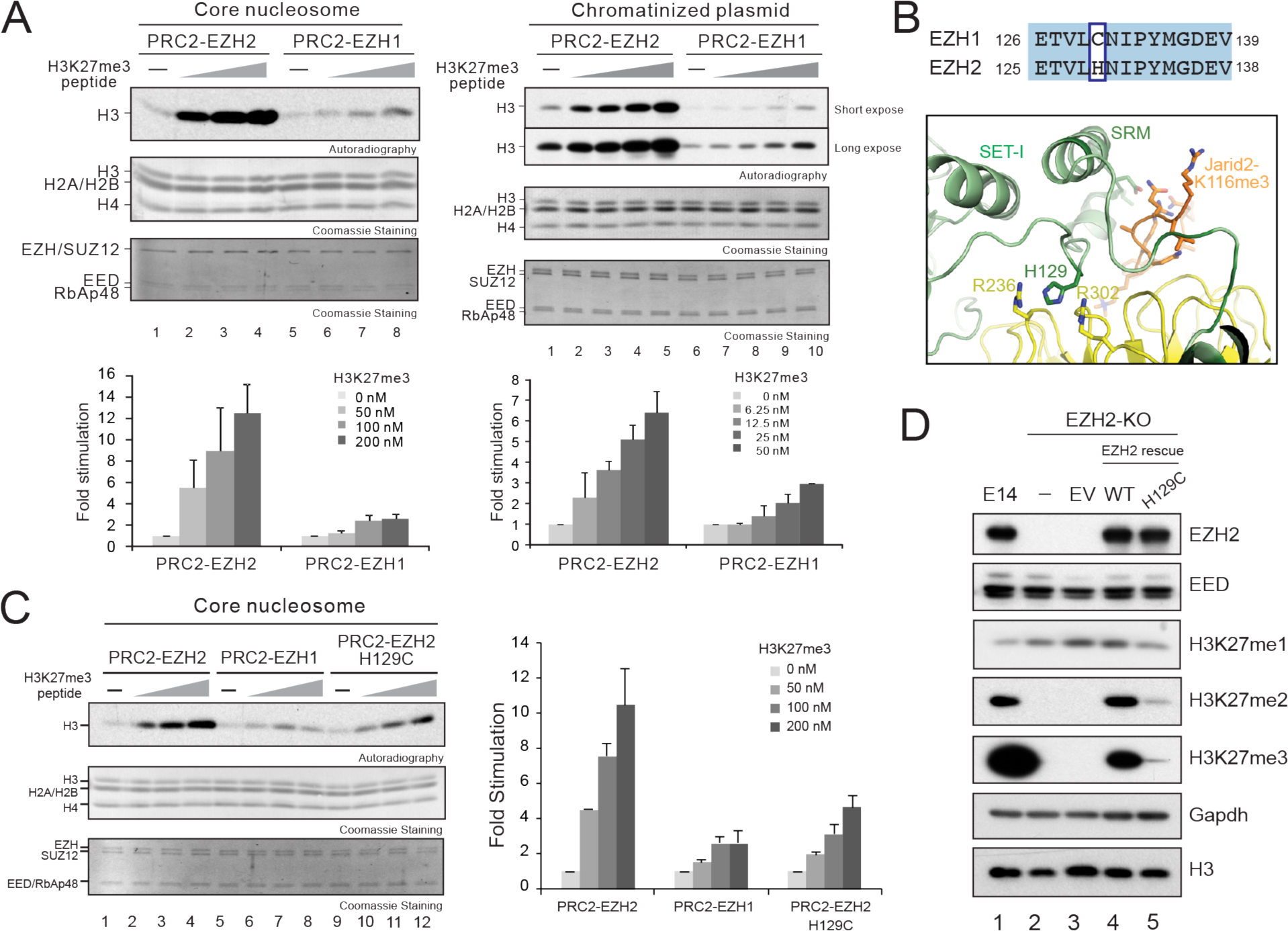
Allosteric, SRM-dependent activation of PRC2-EZH2 is significantly more efficient than that of PRC2-EZH1. (A) *Top*, HMT assays containing PRC2-EZH1 or PRC2-EZH2 (15 nM) using core nucleosomes or chromatinized plasmids as substrates (300 nM) in the absence or indicated amounts of H3K27me3 peptide as illustrated on the panel. Details of the HMT assay conditions are described in the Materials & Methods. The levels of methylation on histone H3 are shown by autoradiography *(Top* image). Coomassie blue staining of SDS-PAGE gels containing nucleosomes *(Middle* image) or PRC2 components *(Bottom* image) was used to visualize the relative concentration of each component present in each reaction. *Bottom*, quantification of the relative amount of ^3^H-SAM incorporated into histone H3 after 60 minutes of incubation (n=3 for each data point). (B) *Top*, Sequence alignment of the SRM domain of human EZH1 and human EZH2. Distinct residues between EZH1 and EZH2 (EZH1 C130 and EZH2 H129) are highlighted by blue box. *Bottom*, Structural depiction of the SRM domain of human PRC2-EZH2 with Jarid2-K116me3 (111-121) peptide. His129 of EZH2 and Arg236 or Arg302 of EED are highlighted. (modified from PDB:5HYN) (C) *Left*, HMT assays containing PRC2-EZH2, PRC2-EZH1, or PRC2-EZH2^H129C^ (15 nM) using core nucleosomes (300 nM) as the substrate in the absence or presence of the H3K27me3 peptide (50, 100, or 200 nM) as illustrated on the panel. Other panels are as described in (A). *Right*, quantification of the relative amount of ^3^H-SAM incorporated into histone H3 after 60 minutes of incubation (n=3 for each data point). (D) Western blot analysis for EZH2, EED, H3K27me1, H3K27me2, H3K27me3, Gapdh, and total histone H3 levels from E14 mESCs WT cells or cells carrying EZH2-KO rescued with EZH2^WT^ or EZH2^H129C^ mutant as indicated. EV, empty vector.

To investigate the basis of this difference, we aligned the sequences of the SRM domains of EZH1 and EZH2, and found one residue (EZH2 H129 and EZH1 C130) that might influence the allosteric activation of PRC2 (Figure 2B, *Top).* Based on the crystal structure of human PRC2 with Jarid2-K116me3 peptide (Justin et al., 2016), EZH2 H129 is positioned between EED residues R236 and R302, and could potentially contribute to the architecture of this region and to the stabilization of the SRM/SET-I interaction resulting in enhanced HMT activity (Figure 2B, *Bottom).* Following this logic, we hypothesized that C130 in EZH1 results in altered interactions in this region, thereby reducing the effectiveness of allosteric activation. To directly test this possibility, we generated EZH2 harboring an H129C mutation (EZH2^H129C^) to mimic EZH1, purified the recombinant PRC2-EZH2^H129C^ complex, and measured its HMT activity. As expected, the allosteric activation of PRC2-EZH2^H129C^ was significantly reduced relative to that of PRC2-EZH2 (Figure 2C). To further explore the differential response to allosteric activation between PRC2-EZH1 and PRC2-EZH2 *in vivo*, we rescued EZH2-KO mESCs with either EZH2^WT^ or EZH2^H129C^. While ectopic expression of EZH2^WT^ rescued H3K27me2-3 levels, ectopic expression of EZH2^H129C^ exhibited marginal rescue effects (Figure 2D, lanes 4 and 5, respectively). Thus, this single residue difference in the SRM domains of EZH1 and EZH2 significantly influenced their catalytic activity in the context of PRC2 *in vitro* and *in vivo.* In addition, our data revealed that allosteric activation of PRC2-EZH2 is necessary for efficient di-and tri-methylation of H3K27 in mESCs. This result is consistent with allosteric activation-deficient mutants showing a complete depletion of H3K27me3 and significantly reduced H3K27me2, while not affecting H3K27me1 in mESCs (Lee et al., 2017).

### PRC2-EZH2 is more responsive to EED or SAM-based inhibitors than PRC2-EZH1

Since allosteric activation of PRC2-EZH2 is more efficient than that of PRC2-EZH1, we reasoned that a small molecule inhibitor that binds to the EED aromatic cage (EED-226) and competes with H3K27me3 would more effectively inhibit PRC2-EZH2. As predicted, increasing amounts of EED-226 compound inhibited the stimulated activity of PRC2-EZH2 and PRC2-EZH1, however, the inhibition was more robust in the case of PRC2-EZH2 (Figure 3A). Next, we treated EZH1^−/−^ or EZH2^−/−^ mESCs with EED-226 for 3 days and monitored the resultant methyltransferase activity. In EZH1-KO cells, H3K27me3 was significantly reduced at all EED-226 concentrations assayed, while H3K27me2 levels were affected modestly (Figure 3B, *Left).* Interestingly, H3K27 mono-methylation accumulated at higher concentrations of EED-226, possibly due to a lack of inhibition of EZH2-mediated conversion to di-and tri-methylated species (Figure 3B, *Left*). These data demonstrated that EED-226 inhibited higher order methylations, i.e. H3K27me2 and-me3, and is less influential on H3K27 monomethylation. In accordance, the level of H3K27 mono-methylation in EZH2-KO cells, devoid of H3K27me2 and me3, was not effectively reduced by EED-226 (Figure 3B, *Right).* Together, these data strongly suggested that in ES cells, PRC2-EZH2 is more responsive to EED-226 than PRC2-EZH1 both *in vitro* and *in vivo.*

**Figure 3.**
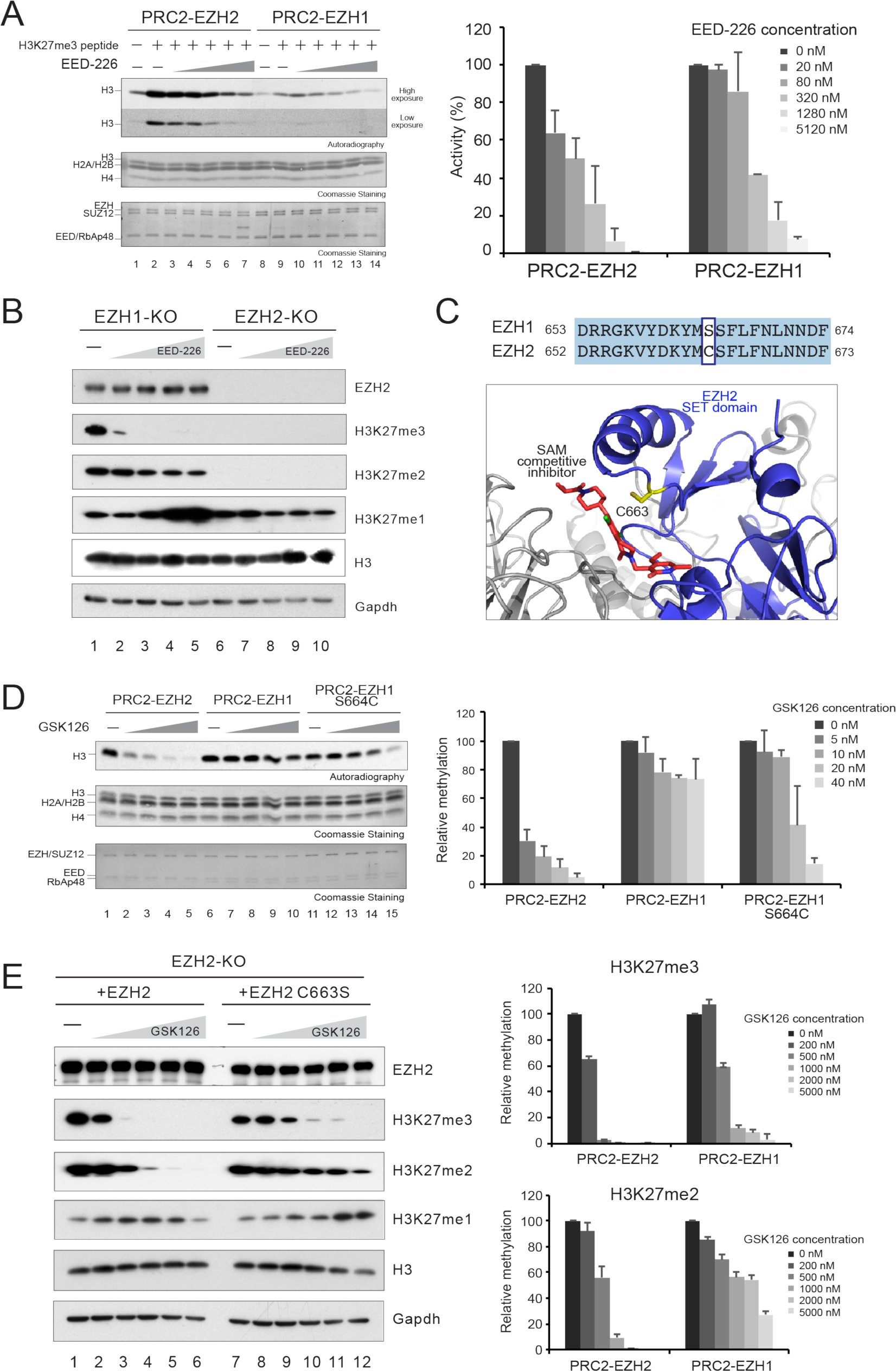
EED inhibitor and GSK126 are more efficient in inhibiting PRC2-EZH2 complex than PRC2-EZH1. (A) *Left*, HMT assays containing 20 nM PRC2-EZH1 or PRC2-EZH2, in the absence or presence of H3K27me3 peptide (300 nM), and increasing amounts (20, 80, 320, 1280, or 5120 nM) of EED inhibitor (EED-226) using di-nucleosomes containing 40 bp linker DNA (300 nM) as the substrate. Details of the HMT assay conditions are described in the Materials & Methods. The levels of methylation on histone H3 are shown by autoradiography *(Top* image). Coomassie blue staining of SDS-PAGE gels containing nucleosomes *(Middle* image) or PRC2 components *(Bottom* image) was used to visualize the relative concentration of each component present in each reaction. *Right*, Relative activity was quantified by the relative amount of ^3^H-SAM incorporated into histone H3 after 60 minutes of incubation (n=3 for each data point). (B) Western blot analysis of EZH2, H3K27me1, H3K27me2, H3K27me3, Gapdh, and total histone H3 levels from EZH2-KO and EZH1-KO. EZH2-KO and EZH1-KO mESCs were treated with increasing amounts of EED-226 (0, 100, 200, 500, or 1000 nM) for 3 days. Whole cell extract was prepared as described in Materials and Methods. (C) *Top*, Sequence alignment of the SET domain of human EZH1 and human EZH2. Distinct residues between EZH1 and EZH2 (EZH1 S664 and EZH2 C663) are highlighted by blue box. *Bottom*, Structural depiction of the SET domain (blue) of human PRC2-EZH2 with an SAM-competitive inhibitor (red). EZH2 C663 residue is highlighted in yellow. (modified from PDB:5IJ7) (D) *Left*, HMT assays containing PRC2-EZH2, PRC2-EZH1, or PRC2-EZH1^S664C^ (15 nM) in the absence or presence of GSK126 (5, 10, 20, or 40 nM) using core nucleosomes (300 nM) as the substrate. Other panels are as described in (A). (E) *Left*, Western blot analysis of EZH2, H3K27me1, H3K27me2, H3K27me3, Gapdh, and total histone H3 levels from E14 mESCs carrying EZH2-KO expressing EZH2^WT^ or EZH2^C663S^. EZH2-KO expressing EZH2^WT^ or EZH2^C663S^ mESCs were treated with increasing amounts of GSK126 (0, 200, 500, 1000, 2000 or 5000 nM) for 3 days. Whole cell extract was prepared as described in Materials and Methods. *Right*, quantification of the levels of H3K27me3 *(Top)* and H3K27me2 *(Bottom)* (n=2 for each data point).

Similar to EED-226, SAM competitive inhibitors are specifically responsive to EZH2 over EZH1. The structure of PRC2-EZH2 with a SAM competitive inhibitor showed that the inhibitor and the SAM binding sites partially overlap (Brooun et al., 2016). One residue (EZH1 S664 and EZH2 C663) close to the SAM binding site was distinctive between EZH1 and EZH2 (Figure 3C). It was suggested that C663 makes van der Waal’s contacts with the bound inhibitor and was predicted to be a key selectivity determinant with respect to EZH1(Vaswani et al., 2016). To test this possibility, we purified PRC2-EZH1^S664C^, which is expected to mimic EZH2, and compared the responsiveness to the SAM competitor inhibitor GSK126. Consistent with previous studies, GSK126 efficiently inhibited the HMT activity of PRC2-EZH2 while only moderately impacting PRC2-EZH1 (Figure 3D). As predicted, PRC2-EZH1^S664C^ showed an increased responsiveness to GSK126 relative to PRC2-EZH1 (Figure 3D). We then introduced EZH2^WT^ or EZH2^C663S^, expected to mimic EZH1, in EZH2-KO mESCs and checked the cellular responsiveness to GSK126. As predicted, H3K27me2/3 was less sensitive to GSK126 in cells rescued with EZH2^C664S^ (Figure 3E). Together, the differences between EZH1 and EZH2 in their SRM and SET domains account for the preferential responsiveness of PRC2-EZH2 to EED-226 and GSK126 inhibitors.

### AEBP2 stimulates PRC2 using a mechanism independent of allosteric activation by H3K27me3

The significant reduction of EZH2 expression during cell differentiation points to EZH1 having a significant role in most differentiated cells (Shen et al., 2008; Son et al., 2013; Stojic et al., 2011). We thus analyzed whether a specific PRC2-related factor, AEBP2, whose expression is apparently not altered upon differentiation (DR, unpublished results) can stimulate PRC2-EZH1 activity. Indeed, AEBP2 stimulated PRC2-EZH1 activity and the fold-stimulation was similar to that of PRC2-EZH2 (Figure 4A and S7). We then analyzed whether allosteric activation (mediated by H3K27me3) was necessary for AEBP2 stimulation. For this, we used a derivative of PRC2 that included the EZH2^P132S^ mutation, which we previously demonstrated to abrogate allosteric activation (Figure 4B, 4C and (Lee et al., 2017)). The results demonstrated that abolishing allosteric activation was without effect on AEBP2-mediated PRC2 stimulation (Figure 4D). Further supporting that allosteric-and AEBP2-mediated activation are additive, we found that the addition of H3K27me3 peptide to reactions containing saturating amounts of AEBP2 resulted in further stimulation of PRC2 activity (Figure 4E).

**Figure 4.**
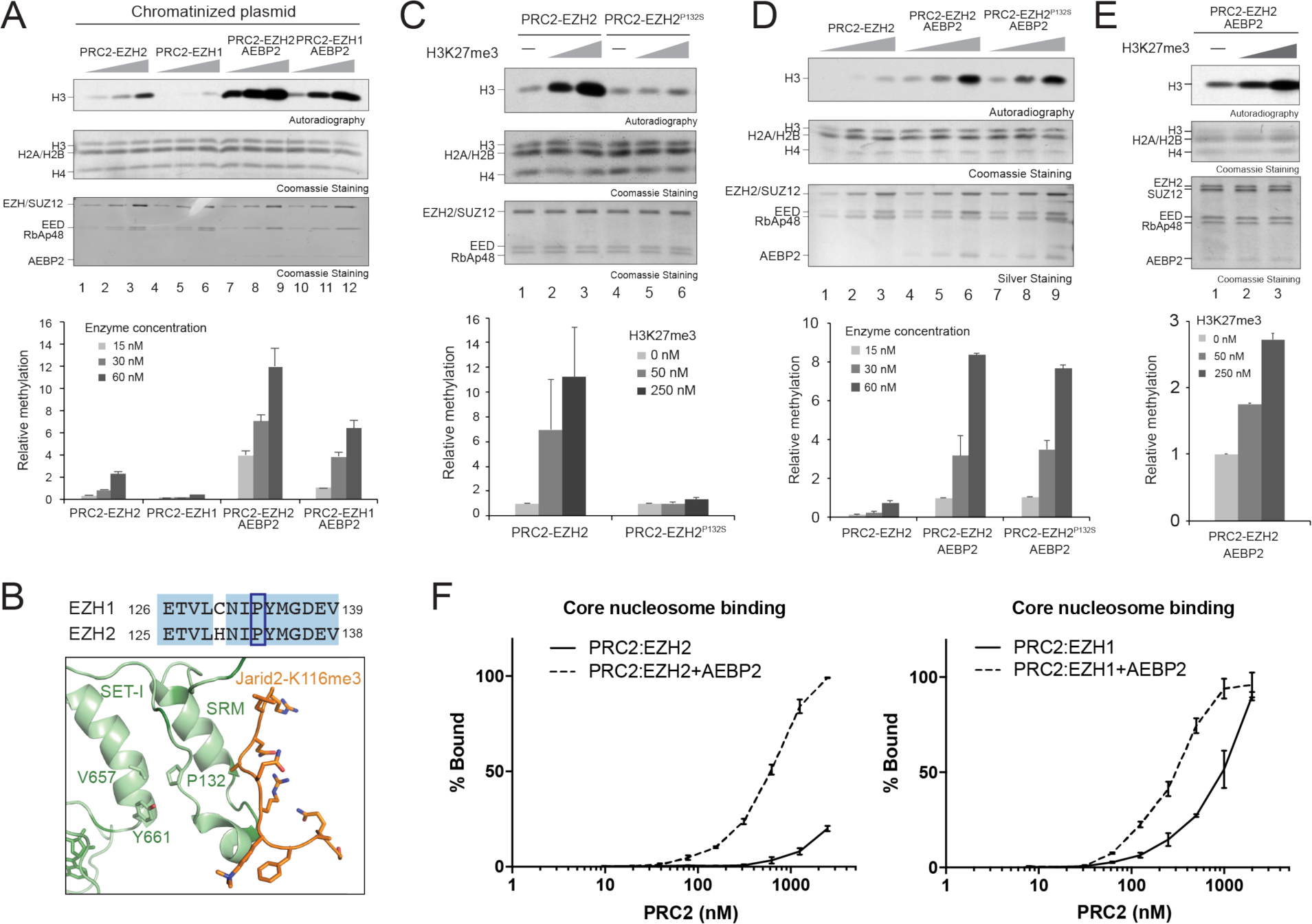
AEBP2 stimulates the HMT activity of PRC2 in an allosteric activation-independent manner. (A) *Top*, HMT assay containing PRC2-EZH1, PRC2-EZH2, PRC2-EZH1-AEBP2, or PRC2-EZH2-AEBP2 (15, 30, or 60 nM) using chromatinized plasmids (300 nM) as the substrate. Details of the HMT assay conditions are described in the Materials & Methods. The levels of methylation on histone H3 are shown by autoradiography *(Top* image). Coomassie blue staining of SDS-PAGE gels containing nucleosomes *(Middle* image) or PRC2 components *(Bottom* image) was used to visualize the relative concentration of each component present in each reaction. *Bottom*, quantification of the relative amount of ^3^H-SAM incorporated into histone H3 after 60 minutes of incubation (n=3 for each data point). (B) Sequence alignment of the SRM domain of PRC2-EZH1 and PRC2-EZH2 (Top). Structural depiction of the SRM domain of PRC2-EZH2 (*Bottom*). (modified from PDB:5HYN) (C) *Top*, HMT assay containing 15 nM PRC2-EZH2 or PRC2-EZH2 containing the EZH2^P132S^ mutant using chromatinized plasmids (300 nM) as the substrate. Other panels are as described in (A). (D) *Top*, HMT assays containing PRC2-EZH2, PRC2-EZH2-AEBP2 or PRC2-EZH2^P132S^-AEBP2 (15, 30, or 60 nM) using chromatinized plasmids (300 nM) as the substrate. Other panels are as described in (A). (E) *Top*, HMT assays containing PRC2-EZH2-AEBP2 (15 nM) with increasing amounts of H3K27me3 peptide (50 or 250 nM) using chromatinized plasmids (300 nM) as the substrate. Other panels are as described in (A). (F) Quantification of EMSA assays containing PRC2-EZH2 *(Left)* or PRC2-EZH1 *(Right)* with or without AEBP2. 200 nM core nucleosomes were used as the substrate. Details of the EMSA assay conditions are described in the Materials & Methods (n=3 for each data point).

Given that the observed AEBP2-mediated stimulation is distinct from the allosteric stimulation, we tested if AEBP2 impacts PRC2 nucleosome binding activity. We performed electrophoretic mobility shift assays (EMSA) using core nucleosomes as the substrate and found that the addition of AEBP2 significantly increased the binding of PRC2-EZH2 (Figure 4F, *Left).* Likewise, AEBP2 stimulated the nucleosome binding of PRC2-EZH1 (Figure 4F, *Right)*, although this stimulation was lower than that of PRC2-EZH2 as PRC2-EZH1 contains higher intrinsic nucleosome binding activity (Son et al., 2013). Taken together, these data indicated that AEBP2 stimulates the HMT activity of PRC2, in part, by enhancing its ability to bind to nucleosomes.

### The KR-motif of AEBP2 is required for enhanced nucleosome binding and PRC2 stimulation

To further investigate the relationship between enhanced nucleosome binding activity and PRC2 stimulation by AEBP2, we used a structure/function approach. Using truncation mutants, we mapped regions of AEBP2 that are important for PRC2 interaction, nucleosome binding, and PRC2 stimulation. First, we determined the PRC2 interaction domain of AEBP2. While AEBP2^(145-295)^, AEBP2^(165-295)^, and AEBP2^(188-295)^ interacted with PRC2, we failed to detect an interaction using AEBP2^(1-187)^ or AEBP2^(1-144)^ (Figure S8). We concluded that the C-terminus of AEBP2, amino acid 188-295, interacts with PRC2. Next, we examined nucleosome binding. The nucleosome binding activity of PRC2-EZH2 containing either full-length AEBP2, AEBP2^(145-295)^ or AEBP2^(165-295)^ was similar, but that of PRC2-EZH2 with AEBP2^(188-295)^ was significantly reduced (Figure 5A and 5B). Therefore, we concluded that the region between amino acids 165 and 188 is important for enhanced nucleosome binding.

**Figure 5.**
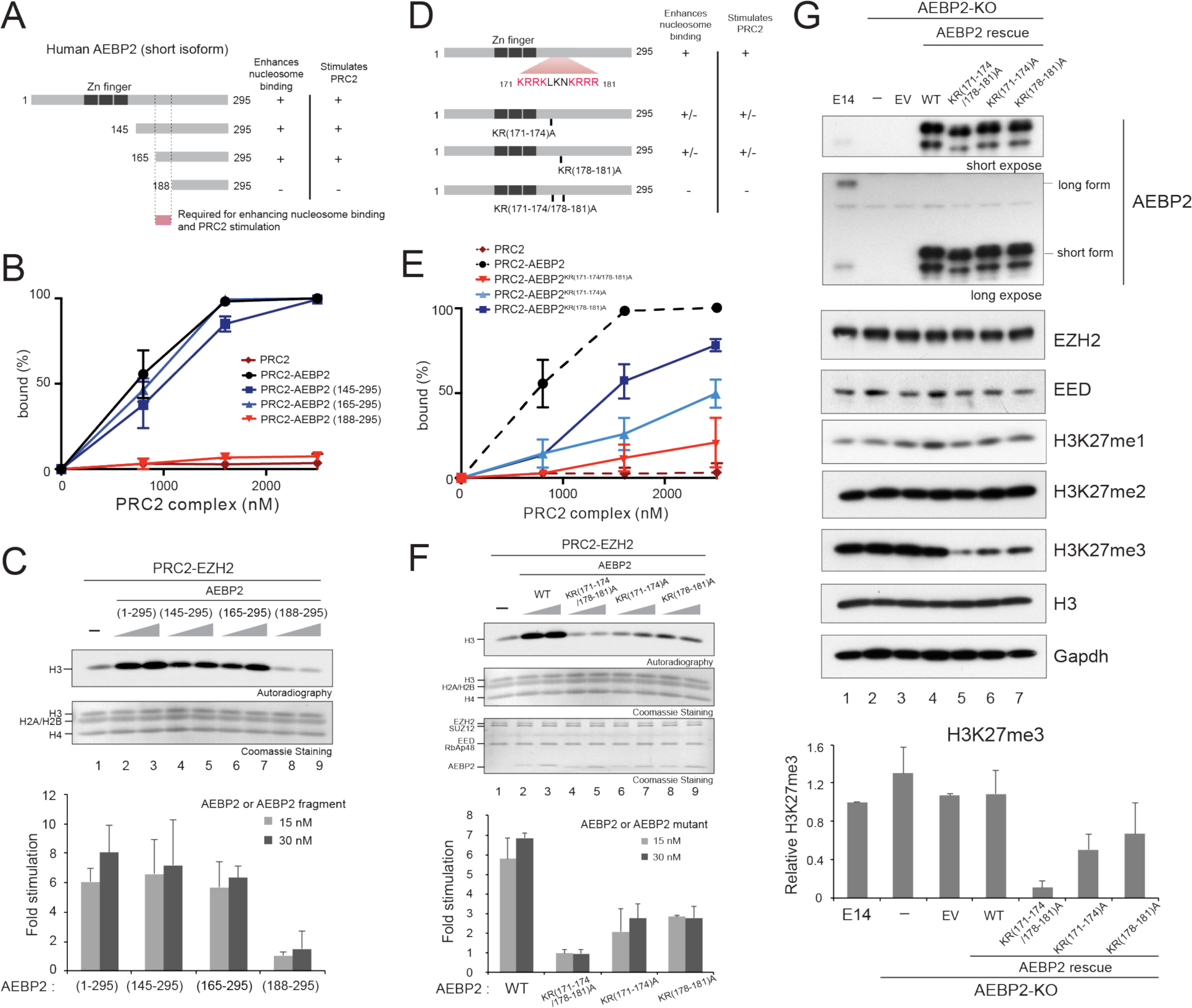
A KR-motif of AEBP2 is required for stimulation of PRC2 catalytic activity. (A) Schematic representation of full-length AEBP2 (short isoform) or its truncated AEBP2 fragments *(Left)*, and the summary of nucleosome binding assays (in Panel B), and HMT assays (in Panel C) for PRC2-EZH2 or PRC2-EZH2 in complex with AEBP2 or AEBP2 fragments. (B) Quantification of EMSA assays containing PRC2-EZH2 or PRC2-EZH2 in complex with AEBP2 or AEBP2 fragments (800, 1600, or 2400 nM) using core nucleosomes as the substrate (200 nM). Details of the EMSA assay conditions are described in the Materials & Methods (n=3 for each data point). (C) *Top*, HMT assays containing PRC2-EZH2 (30 nM) in the absence or presence of AEBP2 or AEBP2 fragments (15 or 30 nM) using core nucleosomes (300 nM) as the substrates. Details of the HMT assay conditions are described in the Materials & Methods. The levels of methylation on histone H3 are shown by autoradiography *(Top* image). Coomassie blue staining of SDS-PAGE gels containing nucleosomes *(Bottom* image) was used to visualize the relative concentration of each component present in each reaction. *Bottom*, quantification of the relative amount of ^3^H-SAM incorporated into histone H3 after 60 minutes of incubation (n=3 for each data point). (D) Schematic representation of AEBP2 or AEBP2 mutants *(Left)* and the summary of nucleosome binding assays (in Panel E), and HMT assays (in Panel F) for PRC2-EZH2 or PRC2-EZH2 in complex with AEBP2, or AEBP2 mutants. (E) Quantification of EMSA assays containing PRC2-EZH2 or PRC2-EZH2 in complex with AEBP2 or AEBP2 mutants (800, 1600, or 2400 nM) using core nucleosomes (200 nM) as substrate (n=3 for each data point). (F) HMT assays containing PRC2-EZH2 (30 nM) in the absence or presence of AEBP2 or AEBP2 mutants (15 or 30 nM) using core nucleosomes (300 nM) as the substrates. Other panels are as described in (C). (G) *Top*, Western blot analysis of AEBP2, EZH2, EED, H3K27me1, H3K27me2, H3K27me3, Gapdh, and total histone H3 levels from E14 mESCs WT cells or cells carrying AEBP2-KO or AEBP2-KO rescued with AEBP2 WT or AEBP2 mutants. *Bottom*, quantification of the levels of H3K27me3, (n=2 for each data point).

Next, we looked at stimulation of methyltransferase activity. Similar to nucleosome binding, the region of AEBP2 that is necessary for PRC2 stimulation mapped between amino acids 165 and 188 (Figure 5C). These data suggested that this region of AEBP2 is required for both enhanced nucleosome binding and PRC2 stimulation. In accordance, AEBP2^(1-187)^ alone interacted with core nucleosomes, while AEBP2^(1-144)^ did not (Figure S9). While inspecting this region of AEBP2, we found a patch of basic amino acids (171-KRRKLKNKRRR-181) that we designated as the KR-motif (Figure 5D). We postulated that this KR-motif is required for both nucleosome binding and PRC2 stimulation. To test this idea, we generated three different clusters of mutations: AEBP2^KR(171-174)A^, AEBP2^KR(178-181)A^, and AEBP2^KR(171-174/178-181)A^ (Figure 5D). We observed a moderate to strong reduction in nucleosome binding in both AEBP2^KR(171-174)A^ and AEBP2^KR(178-181)A^ (Figure 5E, sky blue and dark blue lines, respectively). As expected, we observed a synergistic effect in AEBP2^KR(171-174/178-181)A^, which severely impacted nucleosome binding (Figure 5E, red line). Similar to the impact on nucleosome binding, PRC2-EZH2 with AEBP2^KR(171-174)A^ or AEBP2^KR(178-181)A^ significantly reduced PRC2 stimulation of HMT activity, while PRC2-EZH2 with AEBP2^KR(171-174/178-181)A^ displayed a complete loss of PRC2 stimulation (Figure 5F). This pattern was similar in the case of PRC2-EZH1 (Figure S10). Thus, a basic motif in AEBP2 is required for both nucleosome binding and PRC2 stimulation.

### Mutations in the KR-motif of AEBP2 significantly reduce H3K27me3 *in vivo*

To validate the significance of AEBP2-mediated PRC2 stimulation that we observed *in vitro*, we generated AEBP2-KO mESCs by sequentially targeting both exon 1 and exon 4 using CRISPR-Cas9 technology such that all AEBP2 isoforms were completely depleted (see Materials & Methods for details and Figure 5G, lane 2). As previously reported (Grijzenhout et al., 2016), we did not observe any significant changes in H3K27 methylation in AEBP2-KO by immunoblotting (Figure 5G, lane 2). These cells were then transduced with lentiviruses containing either the wild-type (WT) short form of AEBP2, AEBP2^KR(171-174)A^, AEBP2^KR(178-181)A^, or AEBP2^KR(171-174/178-181)A^, and monitored for changes in H3K27 methylation. We did not observe any significant changes in H3K27 methylation in the WT rescue (Figure 5G, lane 4). In contrast, ectopic expression of AEBP2^KR(171-174)A^ (lane 6) or AEBP2^KR(178-181)A^ (lane 7) displayed reduced H3K27me3, and expression of AEBP2^KR(171-174/178-181)A^ resulted in significantly reduced levels of H3K27me3 (lane 5) (see Discussion). Interestingly, the reduction in H3K27me3 *in vivo* was proportional to the levels of AEBP2-mediated PRC2 stimulation shown in Figure 5F. Neither H3K27me1 nor-me2 were affected (Figure 5G). These data are consistent with AEBP2-mediated PRC2 stimulation being required for maintenance of H3K27me3.

### Multiple means of catalyzing H3K27me3 through PRC2 stimulation

Above, we showed that 40 bp linker DNA and allosteric activation significantly enhanced the catalytic activity of PRC2-EZH2, and that AEBP2-mediated stimulation enhanced both PRC2-EZH1 and PRC2-EZH2 activity. We next tested if each one of these mechanisms can stimulate PRC2 to catalyze the highest extent of H3K27 methylation, H3K27me3, *in vitro.* To this end, we first looked at PRC2 stimulation by an allosteric activator (H3K27me3) or by AEBP2 using H3K27me2-containing nucleosomes and monitored trimethylation of H3K27. Compared to PRC2-EZH2 alone, PRC2-EZH2 with H3K27me3 or AEBP2 displayed significantly enhanced H3K27me3 (Figure 6A, lanes 1-9). The enhanced activity was not observed when using PRC2-EZH2 with the stimulation-deficient AEBP2^KR(171-174/178-181)A^ (Figure 6A, lanes 10-12). Yet, the interpretation of these results with AEBP2 alone might be complicated given that the product of the reaction, H3K27me3, is also an allosteric activator. Thus, we performed a similar experiment using the allosteric deficient mutant, PRC2-EZH2^P132S^. We found that PRC2-EZH2^P132S^-AEBP2 efficiently catalyzed H3K27me3, while PRC2-EZH2^P132S^ alone or PRC2-EZH2^P132S^-AEBP2^KR(171-174/178-181)A^ were ineffectual (Figure 6C, lanes 7-9), suggesting that AEBP2-mediated PRC2 stimulation did efficiently catalyze H3K27me3, and in the absence of allosteric activation. The activity towards tri-methylation of H3K27 was further enhanced when both AEBP2 and H3K27me3 were present, confirming a distinct and additive stimulation (Figure 6A, lanes 13-15). This pattern was similar in the case of PRC2-EZH1, but overall stimulation was much weaker (Figure 6B), highlighting its compromised activity compared to PRC2-EZH2 as well as its compromised response to allosteric activation by H3K27me3.

**Figure 6.**
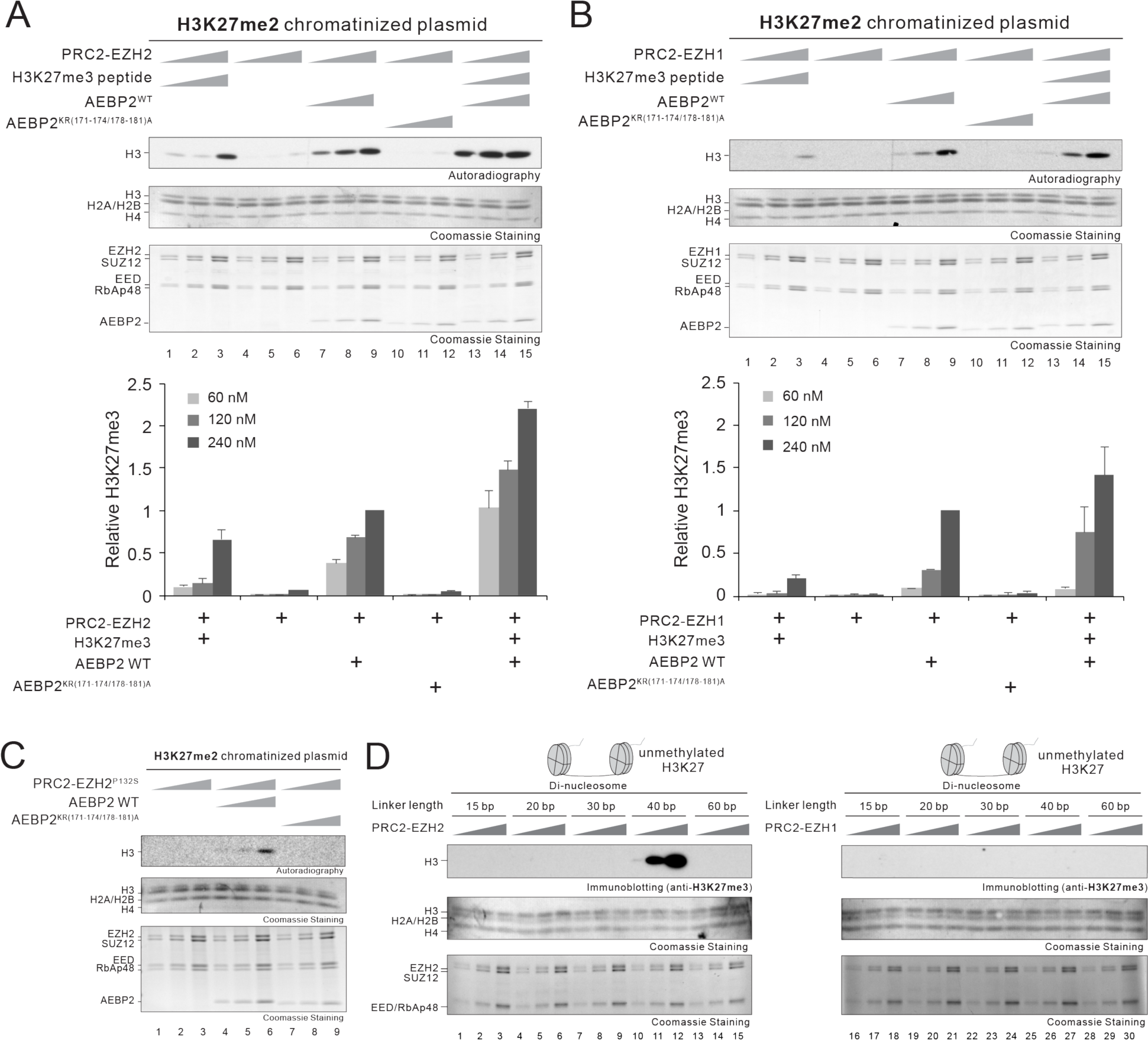
Multiple, independent pathways of catalyzing H3K27me3 through PRC2 stimulation. (A-B) *Top*, HMT assays containing PRC2-EZH2 (A) or PRC2-EZH1 (B) (60, 120, or 240 nM) in the absence or presence of H3K27me3 peptide, AEBP2^WT^, AEBP2^KR(171-174/178-181)A^, or both H3K27me3 and AEBP2^WT^ (60, 120, or 240 nM) using H3K27me2 chromatinized plasmids (300 nM) as substrate. Details of the HMT assay conditions are described in the Materials & Methods. The levels of methylation on histone H3-containing nucleosomal substrates are shown by autoradiography *(Top* image). Note that the signal from autoradiography is the tri-methylation of H3K27. Coomassie blue staining of SDS-PAGE gels containing nucleosomes *(Middle* image) or PRC2 components *(Bottom* image) was used to visualize the relative concentration of each component present in each reaction. *Bottom*, quantification of the relative amount of ^3^H-SAM incorporated (H3K27me3) into histone H3 after 60 minutes of incubation (n=2 for each data point). (C) HMT assays containing PRC2-EZH2^P132S^ (60, 120, or 240 nM) in the absence or presence of AEBP2^WT^, AEBP2^KR(171-174/178-181)A^ (60, 120, or 240 nM) using H3K27me2 chromatinized plasmids (300 nM) as substrate. Other panels are as described in (A-B). (D) HMT assays containing PRC2-EZH2 *(Left)* and PRC2-EZH1 *(Right)* performed in Figure 1D were subjected to immunoblotting against H3K27me3 *(Top* image). Other panels are as described in (A-B).

Next, we checked whether the enhanced activity of PRC2-EZH2 by 40 bp linker DNA, as shown in Figure 1D, was sufficient to catalyze H3K27me3 from unmethylated substrate. Indeed, immunoblotting with antibody against H3K27me3 demonstrated its presence, and notably only in the case of the 40 bp linker DNA (Figure 6D, *Left).* Importantly, and consistent with the results presented above, the stimulatory effect by the 40 bp linker was specific for PRC2-EZH2, as the PRC2 containing EZH1 was not affected (Figure 6D, *Right).* Together, our data revealed multiple and distinct pathways by which PRC2 is stimulated to attain its critical maintenance of H3K27me3 levels.

## Discussion

The diversity amongst histone methyltransferases affords a dynamic regulation of the histone methylation status of chromatin, with consequences to gene expression. In some cases, distinct methyltransferases can perform the same catalysis. For instance, SETD2, NSD1, NSD2, NSD3, and ASH1, are involved in H3K36 methylation, while Suv39h1-2 and G9a/GLP are required for H3K9 methylation (Greer and Shi, 2012). In contrast, PRC2 is recognized as a unique H3K27 methyltransferase in mammals. Indeed, our findings show that cells depleted of both EZH1 and EZH2 activity are devoid of H3K27 methylation (Figure 1A). Yet, as shown here, while the histone methyltransferase activity of core PRC2 is unique, it exhibits stark differences in response to various modulators as a consequence of the identity of the EZH paralog comprising the complex. Such differences reflect an inherent regulation to PRC2 catalytic activity, likely reflecting distinct cellular requirements.

The relative contributions to H3K27 methylation by EZH1 and EZH2 have been extensively studied (Ezhkova et al., 2011; Hidalgo et al., 2012; Margueron et al., 2008; Shen et al., 2008; Son et al., 2013), yet many questions remain regarding their distinct functional roles owing to differences in tissue specific expression (Margueron et al., 2008; Shen et al., 2008; Son et al., 2013). The expression of EZH1 and EZH2 in cells is highly dependent on the stage of cellular differentiation as well as particular cell types (Margueron et al., 2008; Shen et al., 2008; Son et al., 2013). Transcriptome analyses in differentiated cells revealed that most tissues including brain, kidney, prostate, and spleen expressed more EZH1 than EZH2, while tissues such as adipose, parathyroid, and pituitary gland expressed EZH1 exclusively (Shen et al., 2008), suggesting that EZH1 plays important role(s) in postembryonic stages, especially critical in non-dividing cells. Although the expression of EZH1 and EZH2 is comparable in some types of differentiated cells (Shen et al., 2008), EZH2 knockdown in breast cancer cells or EZH2 conditional knockout in neural crest cells significantly reduced H3K27me3 (Gonzalez et al., 2009; Mu et al., 2013; Schwarz et al., 2014), indicating that EZH2 still dominates PRC2 activity in these cases. Hence, not only the expression levels of EZH1 and EZH2 but, more importantly, the regulation of their intrinsic enzymatic properties are critical features of the spatial-temporal regulation of gene repression. Here, we observed that the presence of nucleosome-free DNA, as well as a specific length of linker DNA (40 bp), and the response to allosteric activator significantly influenced and distinguished the catalytic function of these two EZH paralogs. Moreover, AEBP2 stimulated the catalytic function, but irrespectively of the EZH paralog.

One of the notable findings here is that our mechanistic understanding of EZH paralogs reflect and help explain the differential expression and the disparate roles of EZH1 and EZH2 in ES cells versus differentiated cells. In ES cells, the chromatin is more open (Gaspar-Maia et al., 2009, 2011) and has a higher level of H3K27 methylation and therefore, these conditions favor EZH2 that has a higher level of activity (Figure 1B), is less inhibited by free DNA (Figure 1C), and has a greater response to an allosteric activator (Figure 2A). As the H3K27me marks are diluted as cells divide, PRC2-EZH2 is more important for the propagation and maintenance of the H3K27me3 mark. On the other hand, EZH1 is less expressed in ES cells and its activity might be further thwarted by an open chromatin structure due to its strong DNA binding activity (Figure 1C). These features are consistent with PRC2-EZH1 catalyzing predominately monomethylation, while PRC2-EZH2 can catalyze all methylation states in mESCs (Figure 1A). In contrast, in differentiated cells with established H3K27me3-repressed chromatin domains, PRC2-EZH2 is less important and PRC2-EZH1 ‘steps-in’ to maintain these domains. In this case, the chromatin is more compact and PRC2-EZH1 is less inhibited by free DNA. Moreover, PRC2 cofactors such as AEBP2, as reported here, could complement the low intrinsic activity of PRC2-EZH1 (see below).

While it has been shown that chromatin with short nucleosome repeat lengths (NRLs) of 167 bp stimulated PRC2-EZH2 activity (Yuan et al., 2012), we observed that PRC2-EZH2 catalytic activity was significantly increased on chromatin with 187 bp NRLs (Figure 1D and S4), and at least *in vitro*, this increased activity was sufficient to specifically drive H3K27me3 in the absence of any additional factors (Figure 6D). Why would PRC2-EZH2 prefer a specific length of linker DNA? One possibility for the preference is that a proper length of linker DNA spatially positions PRC2 to facilitate catalysis and the establishment of H3K27me3, although a better understanding of the precise manner in which PRC2 is activated by specific NRLs awaits 3D structures of PRC2 with each of its paralogs bound to chromatin. Given the high dependence on particular NRLs for PRC2 activation observed in this study, it is likely that certain ATP-dependent chromatin remodelers cooperate with PRC2 to promote formation of facultative heterochromatin. Similar observations were made for the role of nucleosome remodeling and the deacetylase complex (NuRD), as well as members of the ISWI family in the regulation of heterochromatin (Pegoraro et al., 2009; Postepska-Igielska et al., 2013).

Another important observation here is that PRC2-EZH1 and PRC2-EZH2 respond differently to an allosteric activator (Figure 2A). Moreover, we pinpointed the single residue (H129) in the SRM that is both specific to EZH2 (Figure 2B) and critical for allosteric activation of PRC2-EZH2 (Figure 2C), and most importantly for H3K27me2/3 levels (Figure 2D). Based on the crystal structures of PRC2, H129 can potentially stabilize interactions formed by several residues in EED (R236, H258, and R302) when PRC2-EZH2 is allosterically activated. One possibility is that these interactions are required to anchor the flexible loops of the EZH2-SRM to EED, bringing the SRM helix in proximity to SET-I. Interestingly, three EED residues have been found mutated in Weaver Syndrome patients, all of which are exclusively located at the junction between the EZH2-SRM and EED, suggesting a causal role between allosteric activation of PRC2 and a developmental disorder (Cohen, A.S.A., Gibson, 2016; Cohen et al., 2015; Cooney et al., 2017; Imagawa et al., 2017). Further studies are required to determine the structure of PRC2-EZH1 at a basal as well as activated state. Such structures will facilitate the design of small molecule inhibitors that would either specifically inhibit PRC2-EZH1 or inhibit both PRC2-EZH1/EZH2 complexes at the same time.

Although PRC2-EZH1 is less active than PRC2-EZH2 in actively dividing cells, some tissues are EZH1-dependent and still maintain H3K27me3 levels (Von Schimmelmann et al., 2016). These observations suggest that PRC2 cofactors are critical for complementing the low intrinsic activity of PRC2-EZH1. We demonstrated here that AEBP2 regulates/stimulates PRC2 catalytic activity independently of EZH paralogs. Indeed we found that the KR-motif of AEBP2 is required for PRC2 stimulation. Mutations in this motif significantly reduced the level of H3K27me3 *in vitro* and *in vivo* (Figures 5 and 6). Interestingly, the complete removal of AEBP2 did not affect the levels of H3K27me3 (Figure 5G), likely because other PRC2 partners can bind to PRC2 and compensate for the loss of AEBP2. Indeed, previous findings indicated that MTF2 (PCL2) binding to PRC2 was increased in an AEBP2-KO and that knockdown of PHF1 (PCL1) in HeLa cells led to reduced H3K27me3 levels, suggesting that PCLs could compensate for the absence of AEBP2 (Grijzenhout et al., 2016; Sarma et al., 2008b). On the other hand, compared to the AEBP2-KO, the AEBP2^KR(171-174/178-181)A^ mutant was able to interact with PRC2 likely excluding PCLs from binding to PRC2, yet was deficient in PRC2 stimulation, thereby resulting in a dominant negative effect (Figure 5G). Precisely how the KR-motif stimulates PRC2 activity is not entirely clear, though it appears to be through an enhancement of nucleosome binding (Figure 4F and 5E). In addition to PRC2 stimulation, the enhanced nucleosome binding might also regulate other aspects of PRC2 function such as recruitment to chromatin as well as EZH1-mediated chromatin compaction. Further structural and functional experiments are required to determine these precise roles and mechanisms.

Our mechanistic studies of the EZH paralogs and the PRC2 partner, AEBP2, underscore the distinctive functions of EZH1 and EZH2 in development and the role of AEBP2 as a PRC2 cofactor. In addition, we uncovered the multiple ways by which PRC2 is stimulated to attain higher-order H3K27 methylation. The unique features of these paralogues are underscored by their differential response to specific inhibitors that target either allosteric regulation or direct catalysis by cofactor competition. These findings emphasize that, in addition to PRC2 recruitment, the dynamic regulation of the catalytic activity of PRC2 is critical to understanding its role in maintaining repressed chromatin domains.

## Materials & Methods

### DNA constructs

Baculovirus expression plasmids for human Strep-tagged EZH1 or EZH2, Flag-tagged SUZ12, HIS-tagged EED and HIS-tagged RbAp48 were cloned into pACEBac1 and corresponding donor vectors pIDC, pIDK and pIDS using Gibson Assembly. Vectors were combined using Cre-lox recombination to generate one plasmid encoding all four PRC2 subunits. Human AEBP2, AEBP2 fragments, and mutants were cloned into pGEX-6P-1 vector using Gibson assembly. Wild-type and mutant EZH2 constructs (EZH2^WT^ and EZH2^H129C^, respectively) were subcloned into the pLV-EF1-alpha-IRES-mCherry vector (Clontech).

### Antibodies

Antibodies against EED were produced in-house. Other commercial antibodies against EZH2 (Cell signaling, D2C9, catalog no. 5246), AEBP2 (Cell signaling, D7C6X, catalog No. 14129), H3K27me1 (Millipore, catalog no. 07-448), H3K27me2 (Cell signaling, D18C8, catalog no. 9728), H3K27me3 (Cell signaling, C36B11, catalog no. 9733), H3 (Abcam, catalog no. ab1791), and Gapdh (GeneTex, catalog no. GTX627408) were purchased from the companies indicated.

### Purification of protein using Baculovirus expression system

Recombinant human PRC2 (EZH1 or EZH2, EED, SUZ12, and RbAp48) was produced in High5 (Hi5) insect cells grown in ESF Media (Expression System). After 64 h of infection, Hi5 cells were resuspended in 20 mM Tris-HCl, pH 7.5, 300 mM NaCl, 10% glycerol, and 1 mM DTT supplemented with ROCHE COMPLETE protease inhibitor and lysed with a cell disruptor (550 Sonic dismembrator, Fisher Scentific). The lysate was incubated for 1 h with Strep-Tactin Macroprep beads (IBA). After washing beads with the same buffer, the complex was eluted by proteolytic removal of affinity tags with TEV and 3C protease overnight at 4 °C. Next day, the supernatant was collected and the beads were washed with the same buffer. Fractions were combined and dialyzed against 20 mM Tris-HCl, pH 7.5, 150 mM NaCl, and 1 mM DTT. The complex was further purified by anion-exchange chromatography with a gradient from 0.15-1 M NaCl, followed by size-exclusion chromatography (Superose 6 Increase 10/300GL, GE Healthcare). PRC2-EZH2^P132S^ and PRC2-EZH2^P132S^-AEBP2 were coexpressed in a baculovirus expression system, as described above.

### Bacterial recombinant protein

AEBP2, AEBP2 fragments, and AEBP2 mutants were expressed as N-terminal GST fusions in *E. coli* strain BL21 RIL with LB Media for 16 h at 18 °C by induction with 0.5 mM Isopropyl α-D-1-thiogalactopyranoside (IPTG). *E. coli* cells were resuspended in 20 mM Tris-HCl, pH 7.5, 300 mM NaCl, 10% glycerol, and 1 mM DTT supplemented with protease inhibitor (1 mM phenylmethlysulfonyl fluoride (PMSF), 0.1 mM benzamidine, and 0.625 mg/ml pepstatin A). Salt concentration of the cleared lysate was increased to 1.5 M and DNA was precipitated using 0.15 *%* PEI (pH 7.5). Cleared lysate was applied to GSH Sepharose beads (Macherey-Nagel) for 1 h at 4 °C and eluted by proteolytic removal of the affinity tag with 3C protease, overnight at 4 °C. The supernatant was collected and the beads washed with the same buffer. Fractions were combined and further purified by anion-exchange chromatography with a gradient from 0.3-1 M NaCl, followed by size-exclusion chromatography in 20 mM Tris-HCl, pH 7.5, 300 mM NaCl, and 1 mM DTT (Superdex 75, GE Healthcare).

For PRC2-AEBP2 complex formation, proteins were mixed in a 1:3 ratio (PRC2:AEBP2) in 20 mM Tris-HCl, pH 7.5, 300 mM NaCl, and 1 mM DTT and incubated for 1 h on ice. The complex was subsequently purified by size-exclusion chromatography (Superose 6 Increase 10/300GL column, GE Healthcare).

### Nucleosome reconstitution

Recombinant histones were generated as previously described (Margueron et al., 2008). The methyl-lysine analogue production for the generation of H3K27me2 mimic nucleosomes was previously described in detail (Simon, 2010; Simon et al., 2007). Nucleosomal substrates were reconstituted as described previously (Dyer et al., 2004; Yun et al., 2012).

### HMT assay

Standard HMT assays were performed in a total volume of 15 μl containing HMT buffer (50 mM Tris-HCl, pH 8.5, 5 mM MgCl_2_, and 4 mM DTT) with 500 nM of ^3^H-labeled S-Adenosylmethionine (SAM, Perkin Elmer), 300 nM of nucleosomes, and recombinant human PRC2 complexes under the following conditions. The reaction mixture was incubated for 60 min at 30 °C and stopped by adding 4 μl of STOP buffer (0.2 M Tris-HCl, pH 6.8, 20% glycerol, 10% SDS, 10 mM β-mercaptoethanol, and 0.05% Bromophenol blue). A titration of PRC2 (from 5 to 60 nM) was performed under these conditions in order to optimize the HMT reaction within a linear range, and the yield of each HMT reaction was measured using the following procedures. After HMT reactions, samples were incubated for 5 min at 95 °C and separated on SDS-PAGE gels. The gels were then subjected to Coomassie blue staining for protein visualization or wet transfer of proteins to 0.45 μm PVDF membranes (Millipore) and exposed to autoradiography film (Denville scientific).

### *In vitro* binding assay

Strep-tagged recombinant EZH2 or EZH1 in complex with EED, SUZ12 and RbAp48 were mixed with purified binding partners (8 μg) in 20 mM HEPES, pH 7.5, 50 mM NaCl, 1 mM DTT, 10 % glycerol, and 0.01 % (v/v) Nonidet P-40 in a final volume of 60 μl and incubated for 1 h at 4 °C. Complexes were immobilized on 12 μl Strep-Tactin Macroprep beads, and incubated for another h at 4 °C. The resin was washed three times with 500 μl buffer and eluted with 20 μl of 20 mM HEPES, pH 7.5, 50 mM NaCl, 1 mM DTT, 10% glycerol, and 2.5 mM Destiobiotin and loaded on 15 *%* SDS-PAGE. Proteins were visualized by Coomassie blue staining.

### Gel mobility shift assay

Core nucleosomes or 601 DNA were incubated with PRC2 or PRC2-AEBP2 complexes in 10 mM HEPES, pH 7.9, 50 mM KCl, 1 mM DTT, 0.25 mg/ml BSA, and 5 % glycerol in a total volume of 5 μl for 30 min at room temperature. Complex formation was analyzed by 3.5 % native polyacrylamide gel electrophoreses (0.3 x TBE, 5 % glycerol) followed by Ethidium bromide staining.

### Lentiviral production and delivery

For the production of viral particles, subcloned lentiviral vectors were co-transfected with pcREV, BH-10, and pVSV-G packaging vectors into 293-FT cells. Virus-containing medium was collected 48 h after transfection and polybrene was added to the viral medium at a concentration of 8 μg/ml. The target cells were spin-infected and cells were FACS-sorted for mCherry positive.

### CRISPR/Cas9-mediated genome editing

An optimal gRNA target sequence closest to the genomic target site was chosen using the http://crispr.mit.edu/designtool. The gRNA was cloned into the SpCas9-2A GFP (Addgene: PX458) or SpCas9-2A Puro (Addgene:PX459) via BbsI digestion and insertion(Ran et al., 2013). mESC cells were seeded into 12-well plates at 100,000 cells per well, and transfected with 0.5 μg of the appropriate guide RNAs, template DNA for guide RNAs and Cas9 endonuclease using Lipofectamine 2000 (Life Technologies). The transfection was performed according to the manufacturer’s recommended protocol, using a 3:1 ratio of Lipofectamine:DNA. After transfection, GFP-positive cells were sorted using the Sony SY3200 cell sorter, and 20,000 cells were plated on a 15 cm dish. Puromycin resistant cells were selected in 2 μg/ml puromycin for 48 h and 7-10 days later, single ESC clones were picked, trypsinized in 0.25 % Trypsin-EDTA for 5 min, and plated into individual wells of a 96-well plate for genotyping. Genomic DNA was harvested via QuickExtract (Epicentre) DNA extraction, and genotyping PCRs were performed using primers surrounding the target site. The resulting PCR products were purified and sequenced to determine the presence of a deletion or a mutation event.

EZH1-KO/EZH2^fl/fl^ mESCs were kindly provided by Dr. Tarakhovsky laboratory (von Schimmelmann et al., 2016). pCAGGS-Cre-GFP was transfected into EzH1-KO/EzH2^fl/fl^ mESCs to generate EZH1-KO/EZH2-ΔSET mESCs.

Two sequential deletions were performed to accomplish full knockout of AEBP2. First a gRNA (CCGCGCUCGCCGACAUGGCC) was used to create a frame shift mutation in exon 1 in order to get rid of the long isoform of AEBP2. Second deletion was created using two gRNAs (UUUUGAAGUGAAUUAGUGAA and AUGUUUGUAGAUGCUAAAAG) to delete the entire exon 4 resulting in frame mutation and subsequent loss of short isoform of AEBP2.

## Author contributions

C-H.L., M.H., D.G., D.R., and K-J.A. conceptualized and designed the study. C-H.L., M.H., D.G., R.S-M., R.A.G., J.Z. and M.W. conducted the experiments. M-W.D. made and provided EZH1^−/−^ EZH2^fl/fl^ mESCs. C-H.L., D.G., D.R. and K-J.A. wrote the manuscript.

## Acknowledgement

We thank Drs. L. Vales and J-R.Yu for critical reading of the manuscript as well as past and current Armache and Reinberg members for critical comments and discussion; D. Hernandez and Dr. J-R.Yu for technical assistance; and Dr. A. Tarakhovsky for kindly providing EZH1^−/−^ EZH2^fl/fl^ mESCs. We also thank the NYU Flow Cytometry Core (grant: NIH/NCI P30CA016087) for cell purification. The work in K-J.A.’s laboratory is supported by grants from the Kimmel Scholars Program, the David and Lucile Packard Foundation, and by the National Institutes of Health award (5R01GM115882-03). M.H. was supported by a fellowship from the Alexander von Humboldt Foundation. The work in D.R.’s lab is supported by the National Institutes of Health (NCI R01CA199652) and the Howard Hughes Medical Institute (HHMI).

---

**Figure S1.**
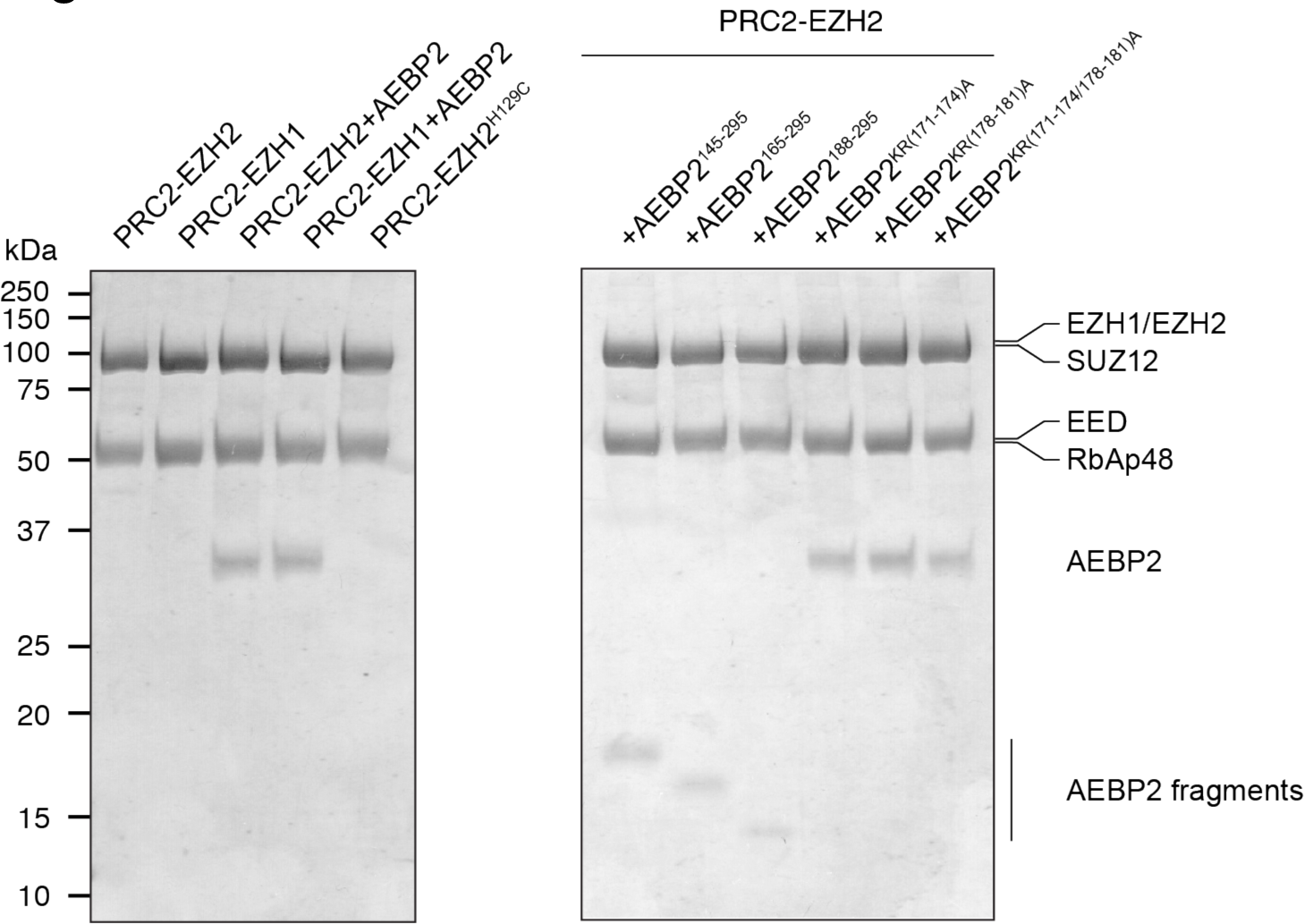
Purification of variant PRC2 complexes. PRC2 complexes were purified as described in Materials & Methods. 8 μg of purified PRC2 complexes were run on a 12 *%* SDS-PAGE gel and stained with Coomassie blue.

**Figure S2.**
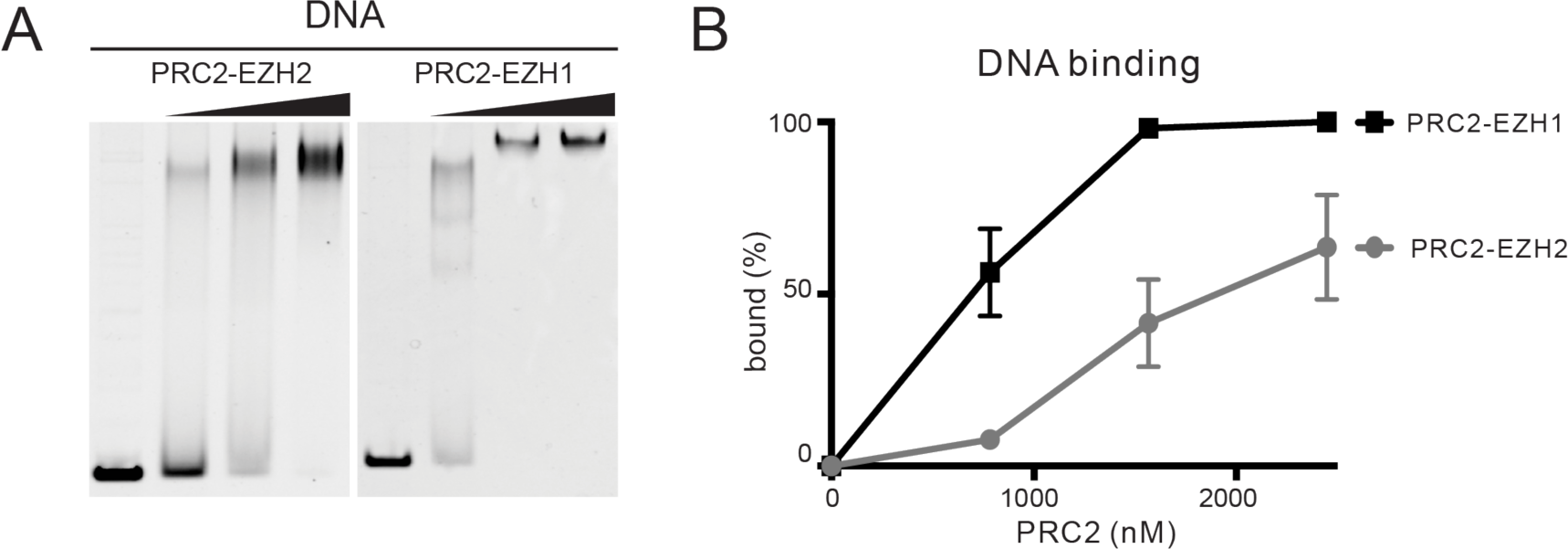
PRC2-EZH1 has higher affinity to DNA than PRC2-EZH2. (A) EMSA assays containing PRC2-EZH2 or PRC2-EZH1 (800, 1600, or 2400 nM) using 147bp 601 DNA (Lowary and Widom, 1998) as the substrate (200 nM). Details of the EMSA assay conditions are described in the Materials & Methods. (B) Quantification of EMSA assays shown in (A) (n=3 for each data point).

**Figure S3.**
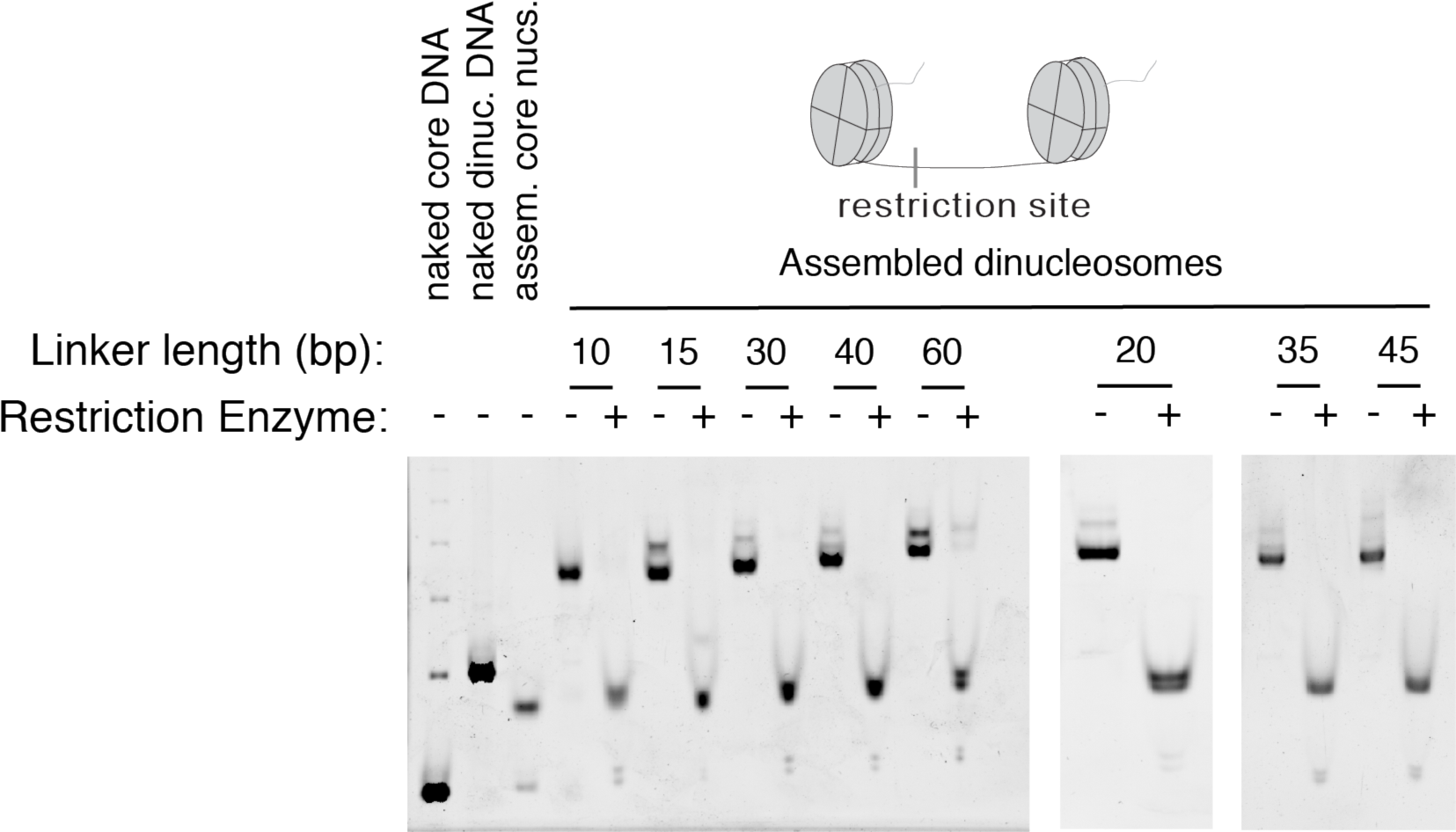
Preparation of di-nucleosomes with variable linker lengths. Di-nucleosome substrates with various length of linker DNAs were treated with or without the indicated restriction enzyme to assess whether the linker remains free of excess octamer. Reactions were run on a 3.5 % native acrylamide gel and stained with ethidium bromide.

**Figure S4.**
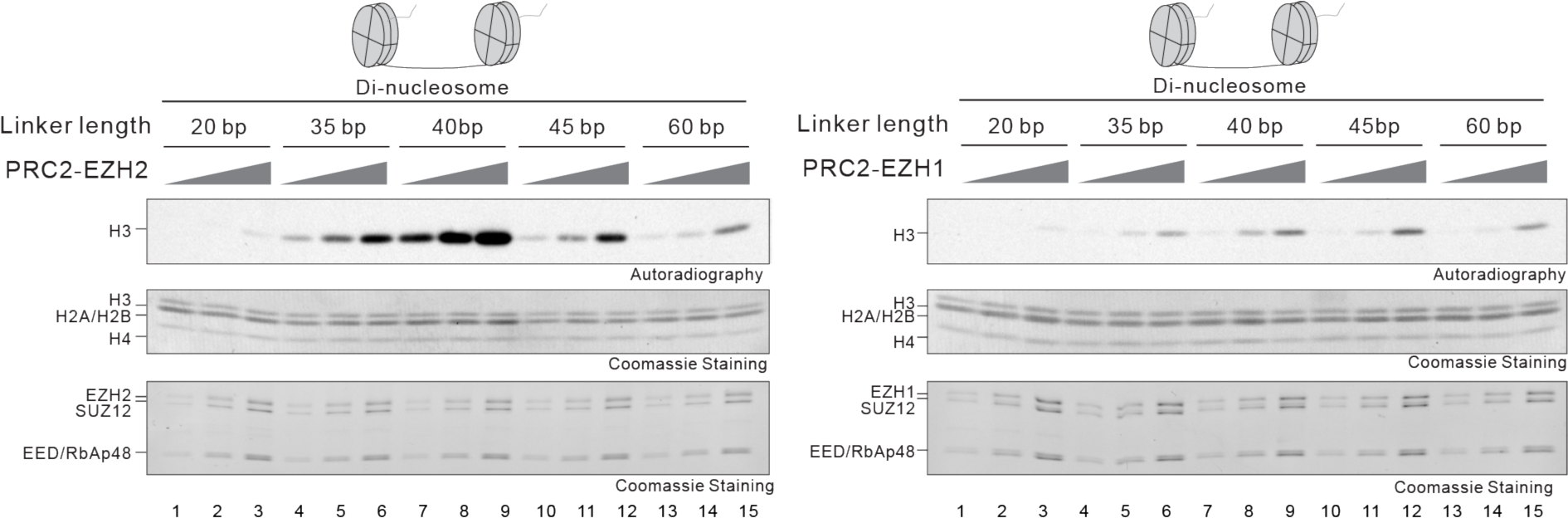
PRC2-EZH2 and PRC2-EZH1 sense nucleosome spacing in a non-linear fashion. HMT assays containing PRC2-EZH1 or PRC2-EZH2 (15, 30, or 60 nM) using di-nucleosomes (300 nM) containing different lengths of linker DNA (20, 35, 40, 45 or 60 bp) as substrates. Note that the flanking DNA at both ends is not present. Details of the HMT assay conditions as well as the generation of the nucleosome substrates are described in the Materials & Methods. The levels of methylation on histone H3 are shown by autoradiography *(Top* image). Coomassie blue staining of SDS-PAGE gels containing nucleosomes *(Middle* image) or PRC2 components *(Bottom* image) was used to visualize the relative concentration of each component present in each reaction.

**Figure S5.**
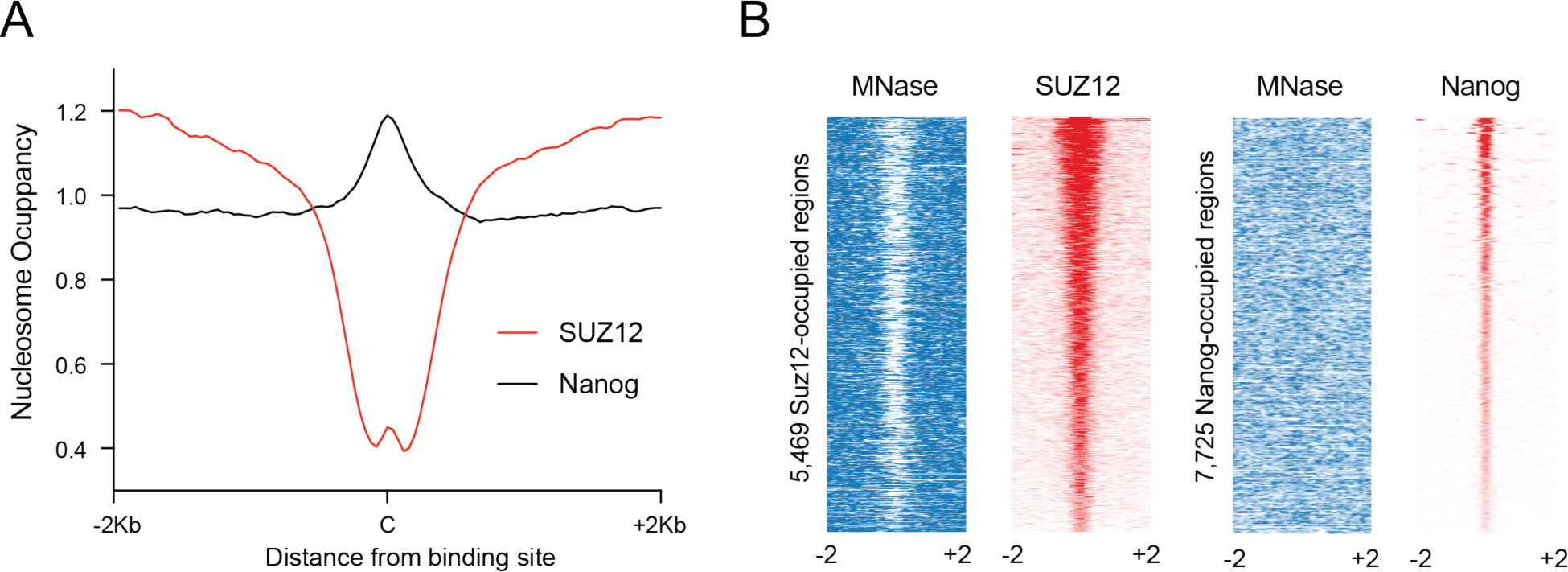
Center of SUZ12 binding sites are devoid of nucleosomes. (A) Genome-wide average nucleosome occupancies at binding sites of SUZ12 and Nanog in ESC. (B) Heatmap representation of ESC MNase-seq (GSE40896) (Teif et al., 2012) and ChIP-seq data at SUZ12 (GSM1397526) (Wei et al., 2015) or Nanog (GSM288345) (Chen et al., 2008)-occupied regions. In all cases, read density is displayed 2 Kb from each side of the center of the binding site.

**Figure S6.**
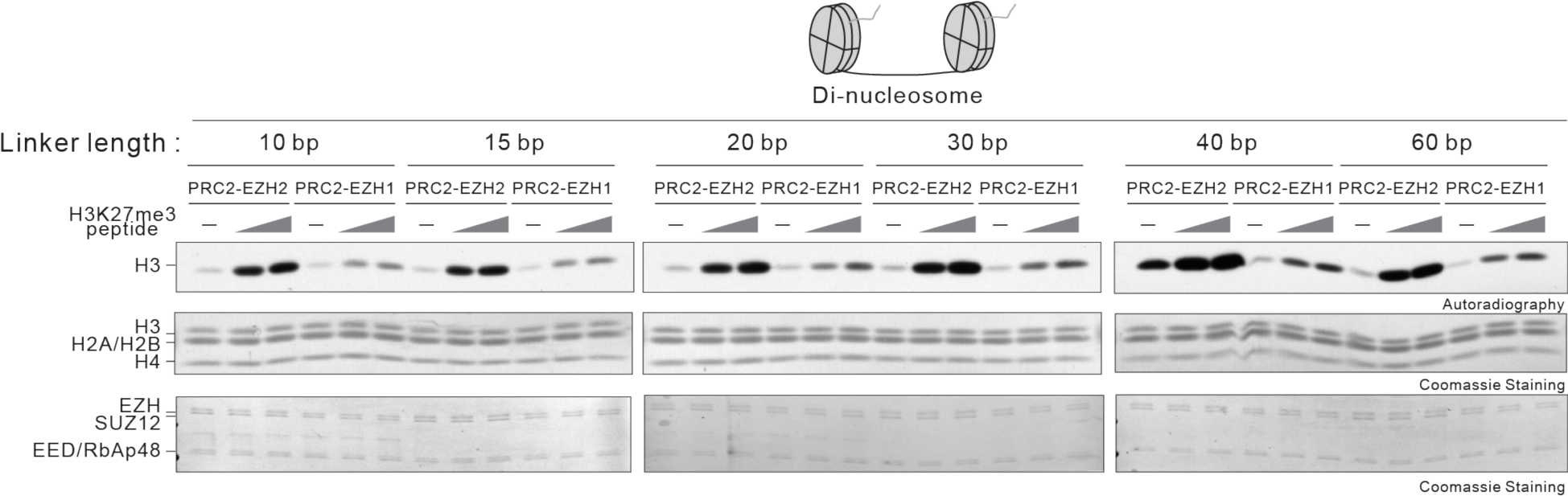
Allosteric activation of PRC2-EZH2 is more efficient than that of PRC2-EZH1. HMT assays containing PRC2-EZH1 or PRC2-EZH2 (15 nM) in the absence or presence of H3K27me3 peptides (50 or 250 nM) using di-nucleosomes (300 nM) containing different length of linker DNA (10, 15, 20, 30, 40, or 60 bp) as substrates. Note that the flanking DNA at both ends is not present. Details of the HMT assay conditions as well as the generation of the nucleosome substrates are described in the Materials & Methods. The levels of methylation on histone H3 are shown by autoradiography *(Top* image). Coomassie blue staining of SDS-PAGE gels containing nucleosomes *(Middle* image) or PRC2 components *(Bottom* image) was used to visualize the relative concentration of each component present in each reaction.

**Figure S7.**
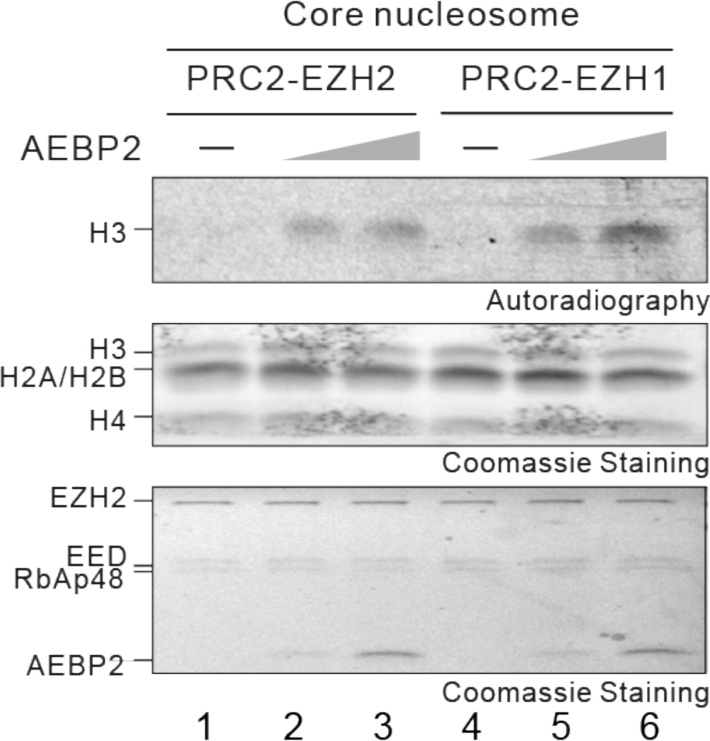
AEBP2 stimulates both PRC2-EZH1 and PRC2-EZH2. HMT assay containing PRC2-EZH1 or PRC2-EZH2 (15 nM) in the absence or presence of AEBP2 (15 or 60 nM) using core nucleosomes (300 nM) as the substrate. Details of the HMT assay conditions are described in the Materials & Methods. The levels of methylation on histone H3 are shown by autoradiography *(Top* image). Coomassie blue staining of SDS-PAGE gels containing nucleosomes *(Middle* image) or PRC2 components *(Bottom* image) was used to visualize the relative concentration of each component present in each reaction.

**Figure S8.**
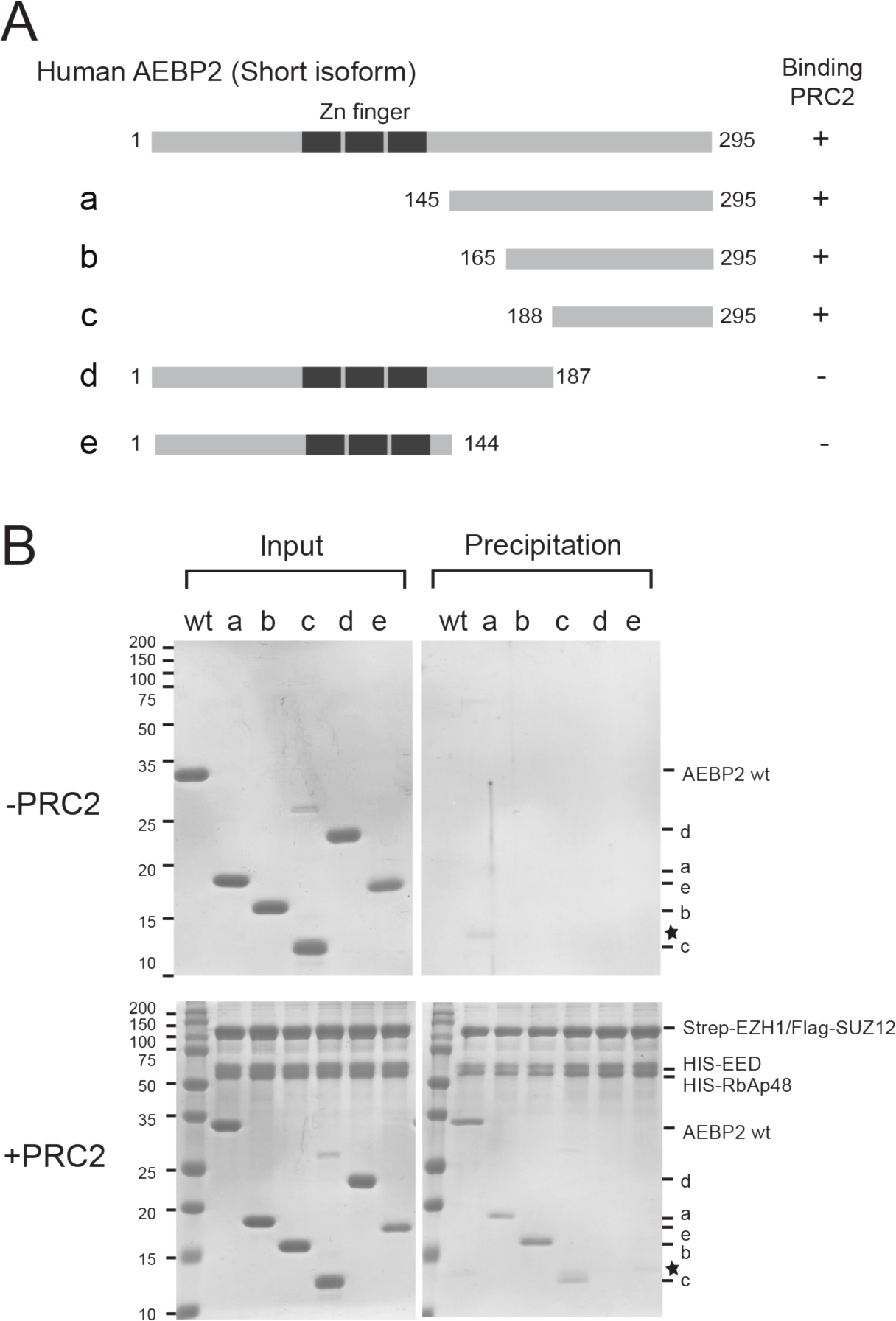
Mapping the PRC2 interaction domain of AEBP2. (A) Schematic representation of AEBP2, AEBP2 fragments *(Left)* and the summary of PRC2 interaction binding results in panel B. (B) AEBP2 or AEBP2 fragments (8 μg) were incubated in the absence *(Top)* or presence (*Bottom*) of 8μg of PRC2 core complex (Strep-EZH1, Flag-SUZ12, 6x HIS-EED, and 6x HIS-RbAp48). Complexes were immunoprecipitated using Streptavidin agarose resin.

**Figure S9.**
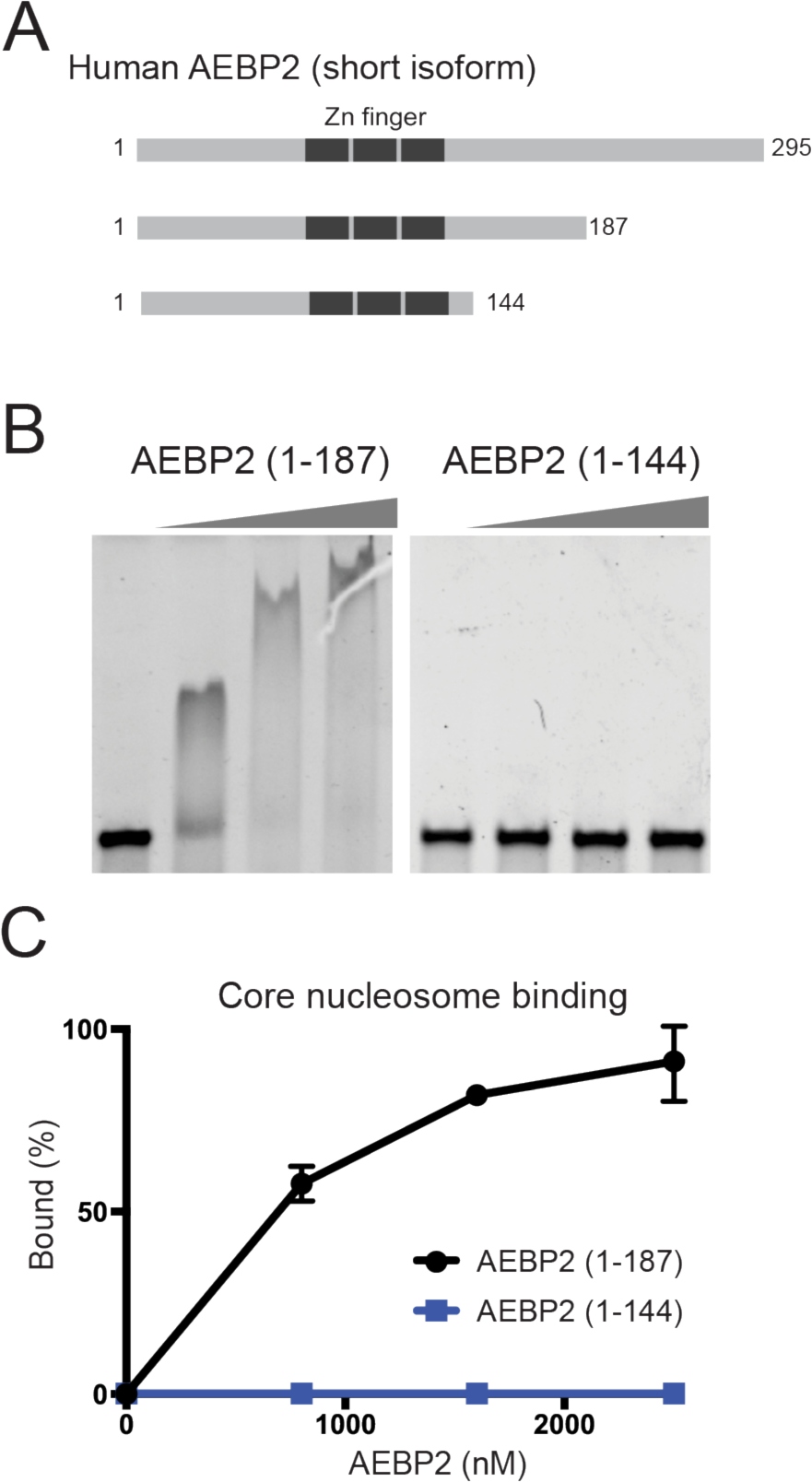
Mapping the nucleosome binding region in AEBP2. (A) Schematic representation of AEBP2 and AEBP2 fragments. (B) EMSA assays containing AEBP2^(1-187)^ or AEBP2^(1-144)^ (800, 1600, or 2400 nM) using core nucleosome as the substrate (200 nM). Details of the EMSA assay conditions are described in the Materials & Methods. (C) Quantification of EMSA assays shown in (A) (n=3 for each data point).

**Figure S10.**
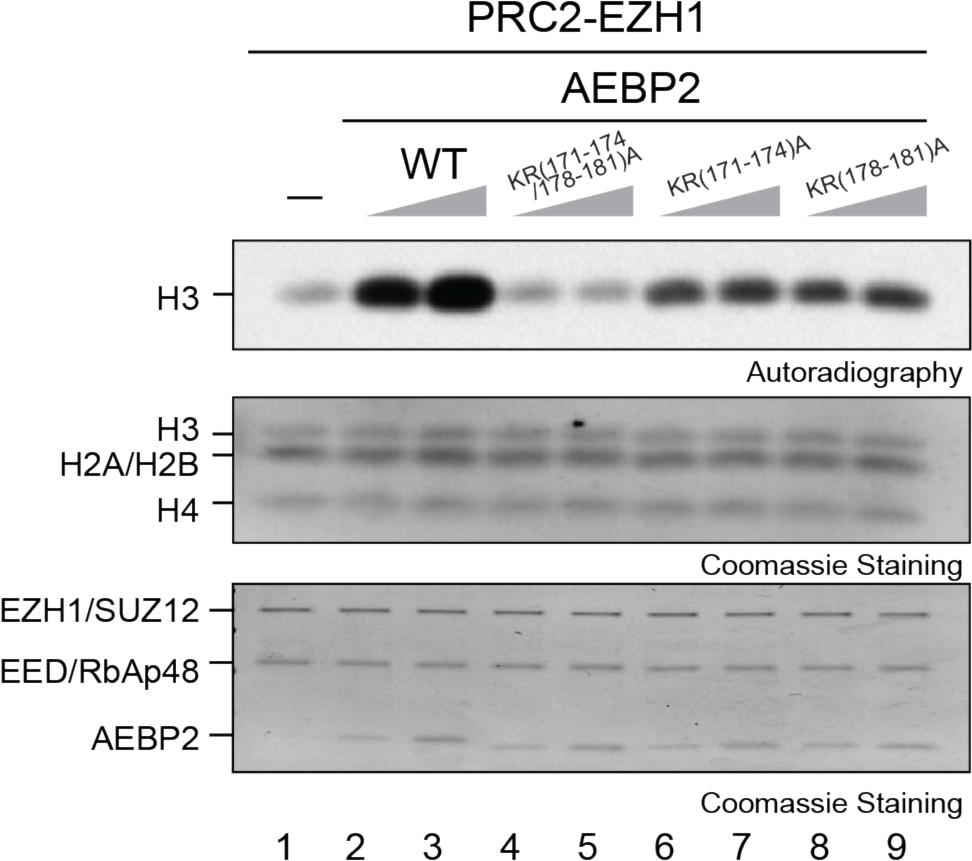
KR-motif of AEBP2 is required for stimulation of PRC2-EZH1. HMT assays containing PRC2-EZH1 (30 nM) in the absence or presence of AEBP2 or AEBP2 mutants (15 or 30 nM) using core nucleosomes (300 nM) as the substrate. Details of the HMT assay conditions are described in the Materials & Methods. The levels of methylation on histone H3 are shown by autoradiography *(Top* image). Coomassie blue staining of SDS-PAGE gels containing nucleosomes *(Bottom* image) was used to visualize the relative concentration of each component present in each reaction.

